# Single-shot 3D photoacoustic tomography using a single-element detector for ultrafast imaging of hemodynamics

**DOI:** 10.1101/2023.03.14.532661

**Authors:** Yide Zhang, Peng Hu, Lei Li, Rui Cao, Anjul Khadria, Konstantin Maslov, Xin Tong, Yushun Zeng, Laiming Jiang, Qifa Zhou, Lihong V. Wang

## Abstract

Imaging hemodynamics is crucial for the diagnosis, treatment, and prevention of vascular diseases. However, current imaging techniques are limited due to the use of ionizing radiation or contrast agents, short penetration depth, or complex and expensive data acquisition systems. Photoacoustic tomography shows promise as a solution to these issues. However, existing photoacoustic tomography methods collect signals either sequentially or through numerous detector elements, leading to either low imaging speed or high system complexity and cost. To address these issues, here we introduce a method to capture a 3D photoacoustic image of vasculature using a single laser pulse and a single-element detector that functions as 6,400 virtual ones. Our method enables ultrafast volumetric imaging of hemodynamics in the human body at up to 1 kHz and requires only a single calibration for different objects and for long-term operations. We demonstrate 3D imaging of hemodynamics at depth in humans and small animals, capturing the variability in blood flow speeds. This concept can inspire other imaging technologies and find applications such as home-care monitoring, biometrics, point-of-care testing, and wearable monitoring.

## Introduction

Vascular diseases, including atherosclerosis, thrombosis, and aneurysms, pose serious health risks such as heart attack, stroke, and organ failure^1^. The early detection of these diseases through imaging of hemodynamics is crucial, enabling prompt intervention and treatment^2^. The efficacy of pharmacological therapies and surgical interventions, such as angioplasty and stenting, can also be evaluated through monitoring changes in blood flow and velocity^3^. Additionally, assessing an individual’s risk of developing vascular diseases is possible by measuring factors such as blood flow velocity, facilitating preventive measures. In biomedical research, imaging of hemodynamics can provide insights into the physiology and pathology of blood vessels, aiding in the development of new treatments for diseases such as hypertension, diabetes, and cancer. Overall, imaging of hemodynamics plays a vital role in disease diagnosis, treatment, and prevention, ultimately improving patient outcomes and advancing medical research^1^.

There are several techniques available to image hemodynamics in the human body, each with its own strengths and limitations. Magnetic resonance imaging (MRI), computed tomography (CT) angiography, and positron emission tomography (PET) are all capable of producing high-resolution images of the vascular system and blood flow dynamics, but they require the use of ionizing radiation and the injection of contrast agents, which can have adverse health effects^4–6^. Moreover, relying on strong ionizing sources and numerous detector elements, these techniques are bulky and expensive, making them inaccessible to mobile clinics or small healthcare facilities. Optical imaging techniques, such as fluorescence imaging and optical coherence tomography (OCT), offer noninvasive visualization of hemodynamics, but their penetration depths are constrained by the optical diffusion limit (~1–2 mm) and do not have sufficient specificity to hemoglobin^7,8^. Doppler ultrasound is another option, providing real-time measurement of blood flow velocity and direction. However, even with recent improvements in minimizing ultrasound probes, state-of-the-art ultrasound imaging techniques still require burdensome and costly data acquisition systems, e.g., Verasonics, due to hundreds to thousands of detector elements^9,10^.

Photoacoustic tomography (PAT), also known as optoacoustic tomography, offers a promising solution to the limitations faced by other imaging techniques for hemodynamic imaging^11–13^.

Unlike other techniques, PAT utilizes the photoacoustic (PA) effect to absorb the energy of incident photons by optical absorbers, such as hemoglobin, in biological tissue and re-emit them as ultrasonic waves (PA waves) to generate optical contrast tomographic images^14–17^. As a result, PAT does not rely on ionizing radiation or contrast agents. Moreover, due to the weak scattering of ultrasound in biological tissue, PAT provides a depth-to-resolution ratio of approximately 200, enabling high spatial resolution at depths up to several centimeters^11^. Two primary forms of PAT are photoacoustic microscopy (PAM)^17–19^, which requires sequential scanning of the probing beam, and photoacoustic computed tomography (PACT)^15,20,21^, which captures a 3D image using one or a few pulses of the probing beam. PAM utilizes a single-element detector, which requires a simple data acquisition system but suffers from low imaging speed. In contrast, PACT offers higher imaging speed but requires numerous detection elements and the corresponding data acquisition systems, making it complex, expensive, and bulky.

To overcome the challenges of complexity, cost, and size in PACT systems, researchers have been working on ways to reduce the number of detector elements needed to reconstruct a 3D image. One promising approach utilizes the principles of compressive sensing and single-pixel imaging^22–25^, which use acoustic scatterers to achieve PA or ultrasound tomography with just a single detector element^26–29^. However, these techniques are time-consuming, as they require a sequence of measurements with different mask configurations, limiting their speed. To address this issue, researchers have developed novel methods that take advantage of the spatiotemporal encoding of an ergodic relay (ER) or a chaotic cavity^30,31^. These techniques can produce high-quality single-shot images while using fewer detector elements^32–36^. However, they have only been demonstrated for 2D imaging and require recalibration for different objects, which can be time-consuming. Additionally, they may not be suitable for long-term imaging in unstable environments due to their sensitivity to boundary conditions.

Here, we present photoacoustic computed tomography through an ergodic relay (PACTER), a PACT system that simultaneously addresses the challenges faced by other imaging techniques. PACTER provides a highly accessible and efficient solution, paving the way for noninvasive, label-free, and ultrafast 3D imaging of hemodynamics at depth in humans. With PACTER, a single-element detector encodes information equivalent to that of 6,400 virtual ones, enabling the reconstruction of a tomographic image of vasculature in 3D with just a single laser pulse. The system achieves ultrafast volumetric imaging at kilohertz rates, making it possible to capture fast hemodynamics in the human body in real-time. We demonstrate PACTER’s capability in monitoring vital signs in small animals and visualizing human hemodynamics in response to cuffing, capturing the variability in blood flow speeds. Because PACTER signals are unaffected by the boundary conditions of the object, the system only needs to be calibrated once and is suitable for long-term imaging in an unstable environment. PACTER’s single-element detector design makes it convenient, affordable, and compact, thus translatable to clinical applications such as home-care monitoring ^37,38^, biometrics, point-of-care testing^39^, and noninvasive hemodynamic monitoring in intensive care units^40^. The single-element detector concept in PACTER can also be generalized to other imaging technologies, such as ultrasonography^41^, sonar^42^, and radar^43^.

## Results

### PACTER system

The PACTER system requires calibration only once prior to its utilization for a complete series of imaging. (Methods). In the calibration procedure (Fig. 1a), the laser beam was transmitted through the ER and focused on a uniform optical absorber placed on top of the ER. We used bovine blood as our calibration target (Supplementary Fig. 1). Using two motorized stages, we controlled the positions of a pair of mirrors to steer the focused laser beam across the field of view (FOV) in the *x-y* plane and recorded the PACTER signals at each scanning position. After calibration, the uniform optical absorber could be removed, and the system was ready for imaging. In the imaging procedure (Fig. 1b), the focusing lens was replaced by a fly’s eye homogenizer (Supplementary Fig. 2), which converted the incident laser beam into a widefield, homogenized illumination pattern that had the same shape and width as that of the calibration FOV (Supplementary Note 1). To acquire imaging data, the object was directly placed on top of the ER using ultrasound gel as the coupling medium, and we recorded the PACTER signal generated by the object following each laser pulse.

**Fig. 1.**
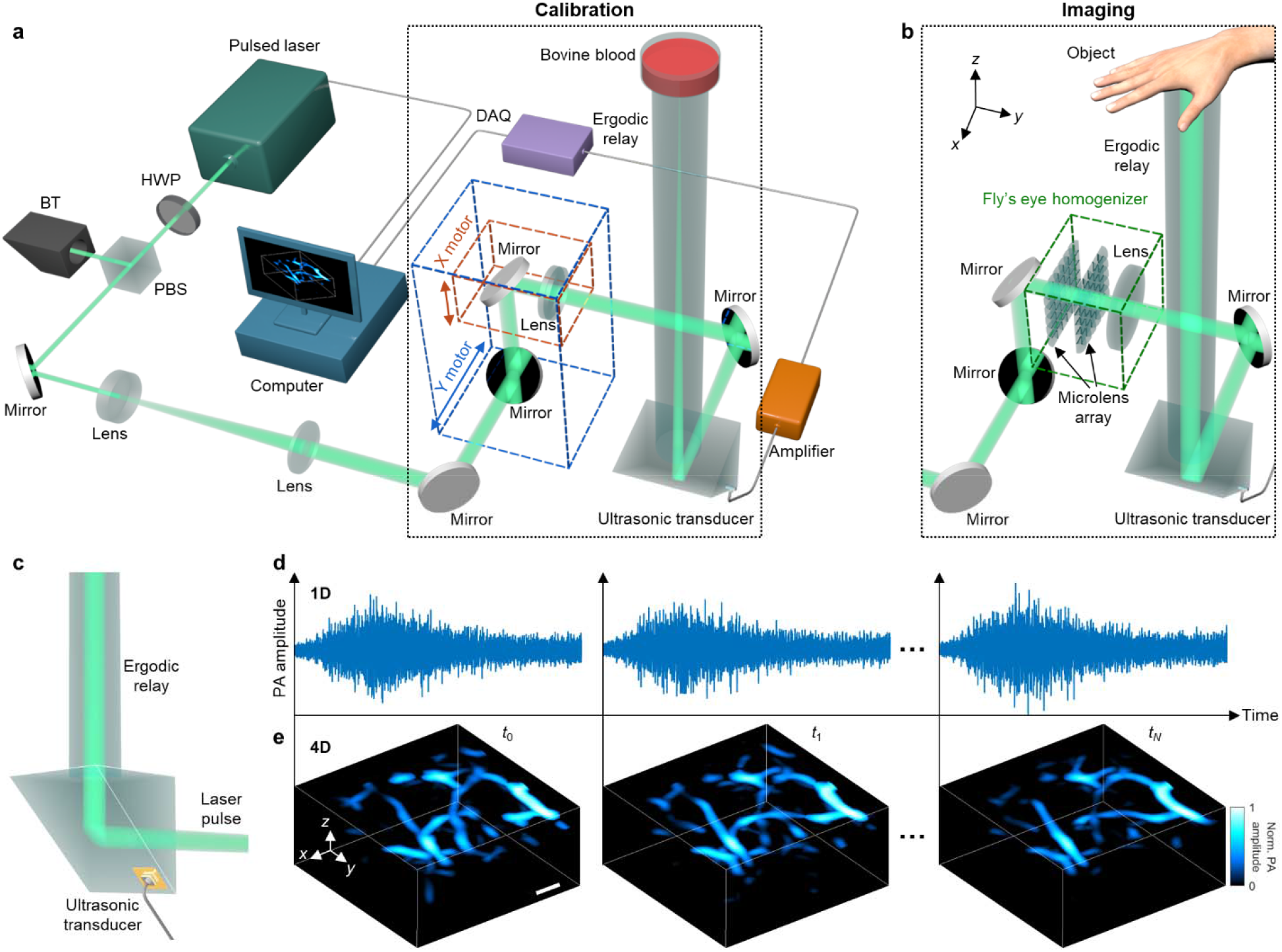
PACTER system. **a,b**, Schematic of the PACTER system in the calibration (**a**) and imaging (**b**) procedures. HWP, half-wave plate; PBS, polarizing beamsplitter; BT, beam trap; DAQ, data acquisition unit. The differences between the two modes are highlighted in the black dotted boxes. **c**, Schematic of the single-element ultrasonic transducer fabricated on the ER. **d**, 1D (*t*) PACTER signals detected by the ultrasonic transducer at time instances *t*_0_, *t*_1_, and *t_N_*. **e**, Reconstruction of a 4D (*xyzt*) image of human palmar vessels from the signals in **d**. Norm., normalized. Scale bar, 1 mm.

To enhance the detection sensitivity and improve the stability of the system, we fabricated a single-element ultrasonic transducer on the hypotenuse surface of the prism of the ER (Fig. 1c).

The transducer was based on a lead magnesium niobate–lead titanate (PMN-PT) single crystal (Supplementary Figs. 3 and 4), which achieved exceptional piezoelectric performance, such as high piezoelectric constant (*d_33_*) and electromechanical coupling coefficient (k_t_)^44–46^. Further, because one of the gold electrodes of the transducer was directly sputtered on the surface of the ER, the PACTER signals inside the ER could reach the transducer with the maximum transmission, and the transducer and the ER became a whole piece that facilitated stable data acquisition.

Conventionally, a single-element ultrasonic transducer can only acquire a 1D signal in the time domain (Fig. 1d). However, with the ER, PACTER can use the single-element transducer to encode spatiotemporal information equivalent to those captured by 6,400 detectors (Supplementary Video 1), which can then be used to reconstruct a 3D map of the optical absorbers in the imaging volume. With a kilohertz laser repetition rate, PACTER can use the 1D (*t*) signals to generate a thousand 3D (*xyz*) volumes per second, leading to a high-speed 4D (*xyzt*) image of optical absorption in, e.g., human palmar vessels (Fig. 1e).

### PACTER signal and reconstruction

PACTER needs only a one-time universal calibration despite its more stringent requirement for 3D imaging (Supplementary Note 2, Supplementary Fig. 5, Supplementary Video 2). In the calibration procedure, the focused laser beam was scanned across the FOV in 80 by 80 steps with a step size of 0.1 mm (Fig. 2a). To ensure that the PACTER signal acquired at each calibration pixel was distinct from others, we chose the step size to be about a half of the full width at half maximum (FWHM) of a line profile along the cross-correlation map, i.e., ~0.21 mm (Supplementary Fig. 6). Although the calibration signals were obtained by scanning the laser beam across a 2D plane, they could be used as 80 × 80 = 6,400 virtual transducers for 3D reconstruction (Fig. 2b) because (1) the PACTER signals were object-independent, and (2) the calibration signals were generated at the bottom of the 3D imaging volume. When source points in the 3D volume 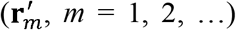 were illuminated by a laser pulse, the PA signals they generated would propagate to the calibrated virtual transducers (*r_n_, n* = 1, 2,…) after time 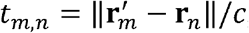, where *c* is the speed of the sound in the medium. Then, these signals would follow the same acoustic path inside the ER to the ultrasonic transducer as that of the calibration signals. From the transducer’s perspective, compared with the calibration signal *k_n_*(*t*) acquired at ***r**_n_* (Fig. 2c), the signal from the source point 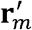 relayed through *r_n_* is proportional to *k_n_*(*t*) delayed by *t_m,n_*, i.e., 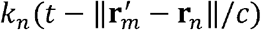 (Fig. 2d). The signal is modulated by both *p*_0,*m*_, the initial pressure at 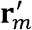, and a weighting factor dependent on the angle and distance. Accordingly, we developed an algorithm to reconstruct the initial pressure in the 3D volume (Supplementary Note 3). Because the reconstruction is prohibitively computationally intensive, we reformulated the forward model through temporal convolution and implemented it using the fast Fourier transformation, increasing the computational efficiency by 48,000 times.

**Fig. 2.**
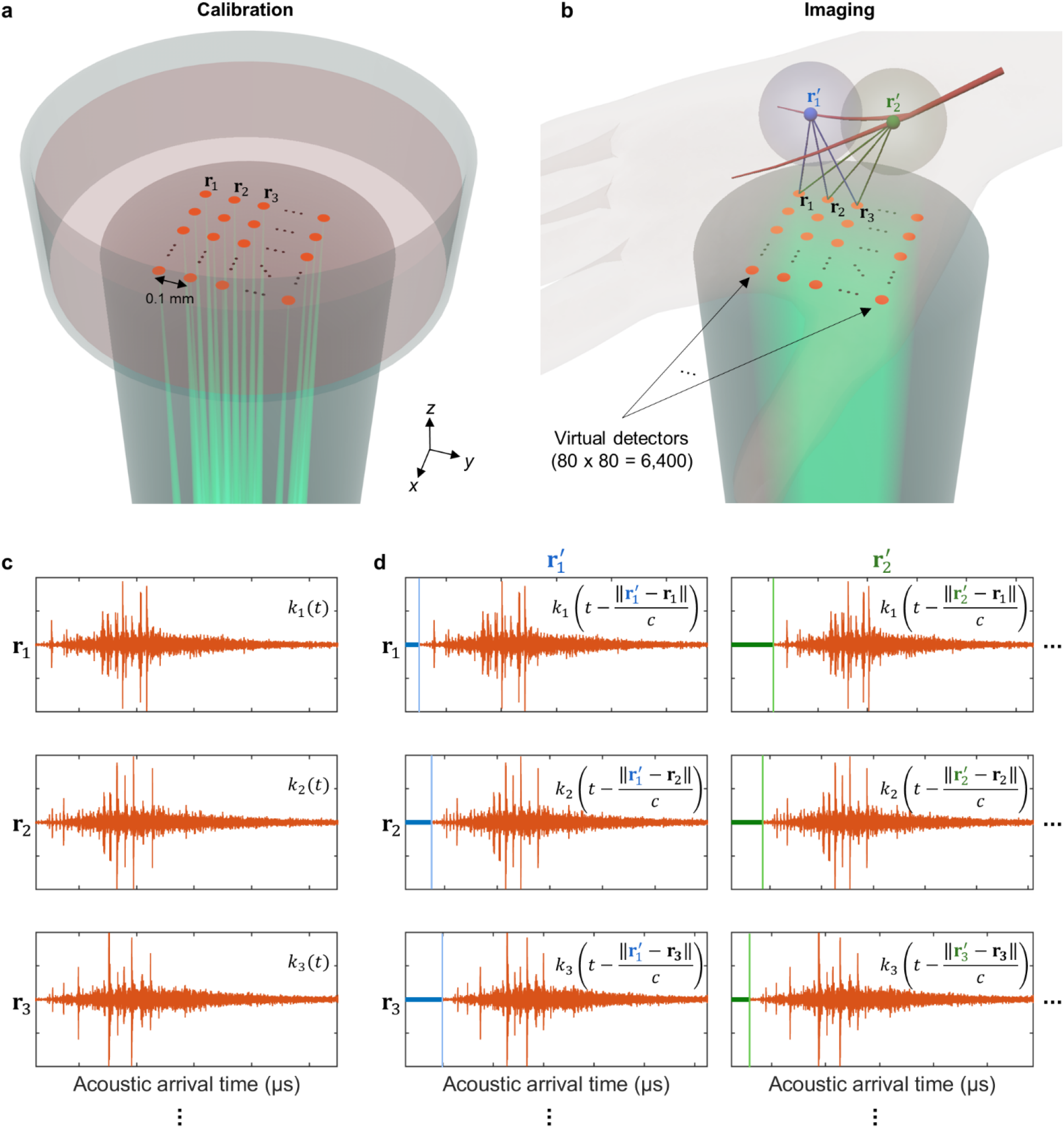
Single-shot 3D reconstruction in PACTER. **a**, Illustration of the calibration procedure of PACTER. Focused laser beams for calibration are shown in green. Calibration pixels are highlighted as orange dots. Calibration step size is 0.1 mm. The calibration pixels (80 x 80) become 6,400 virtual transducers. **r**_1_, **r**_2_, **r**_3_ are the positions of three calibrated virtual transducers. **b**, Illustration of PACTER of human palmar vessels. The homogenized beam for widefield illumination is shown in green. 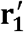 and 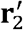 are the positions of two source points in the vessels. Blue and green spheres denote the PA waves generated by the source points. The calibrated virtual transducers capture the PA signals from 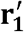 and 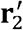 with different delays, indicated by the thick blue and green lines. **c**, PACTER signals, *k*_1_(*t*), *k*_2_(*t*), *k*_3_(*t*), of the calibrated virtual transducers corresponding to **r_1_ r_2_, r_3_**, respectively. **d**, PACTER signal from the widefield imaging consists of PA signals from 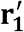 and 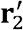, which are essentially *k*_1_(*t*), *k*_2_(*t*), *k*_3_(*t*) delayed according to the distance between the calibrated virtual transducer and the source point.

### Spatiotemporal characterization of PACTER

Using the signals acquired by the single detector, PACTER can image the 3D structure of a curved black wire with a single laser pulse (Fig. 3a) and the 4D dynamics of bovine blood flushing through an S-shaped tube when the tube was illuminated by multiple laser pulses (Fig. 3b, Supplementary Video 3). To evaluate whether the 3D volumes reconstructed by PACTER were correct measurements of the objects, we first compared the perspective views of the 3D volumes reconstructed by PACTER with the ground-truth projection images formed by raster-scanning the laser beam across the objects (Supplementary Fig. 7). Despite a lower spatial resolution compared with the ground truth, the comparison demonstrates that PACTER can correctly reconstruct the 3D objects in the lateral (*x*-*y*) directions. Second, we imaged a thin object in water, whose *z* positions were precisely controlled and measured by a linear translation stage. We imaged the object at multiple *z* positions, reconstructed the 3D volumes (Fig. 3c), and compared the *z* positions in the reconstructed volumes with the real ones. As shown in Fig. 3d, the reconstructed and real *z* positions were linearly related (*R*^2^ = 1.000) with a slope (0.993) close to 1, demonstrating that PACTER can accurately reconstruct the 3D objects in the axial (*z*) direction.

**Fig. 3.**
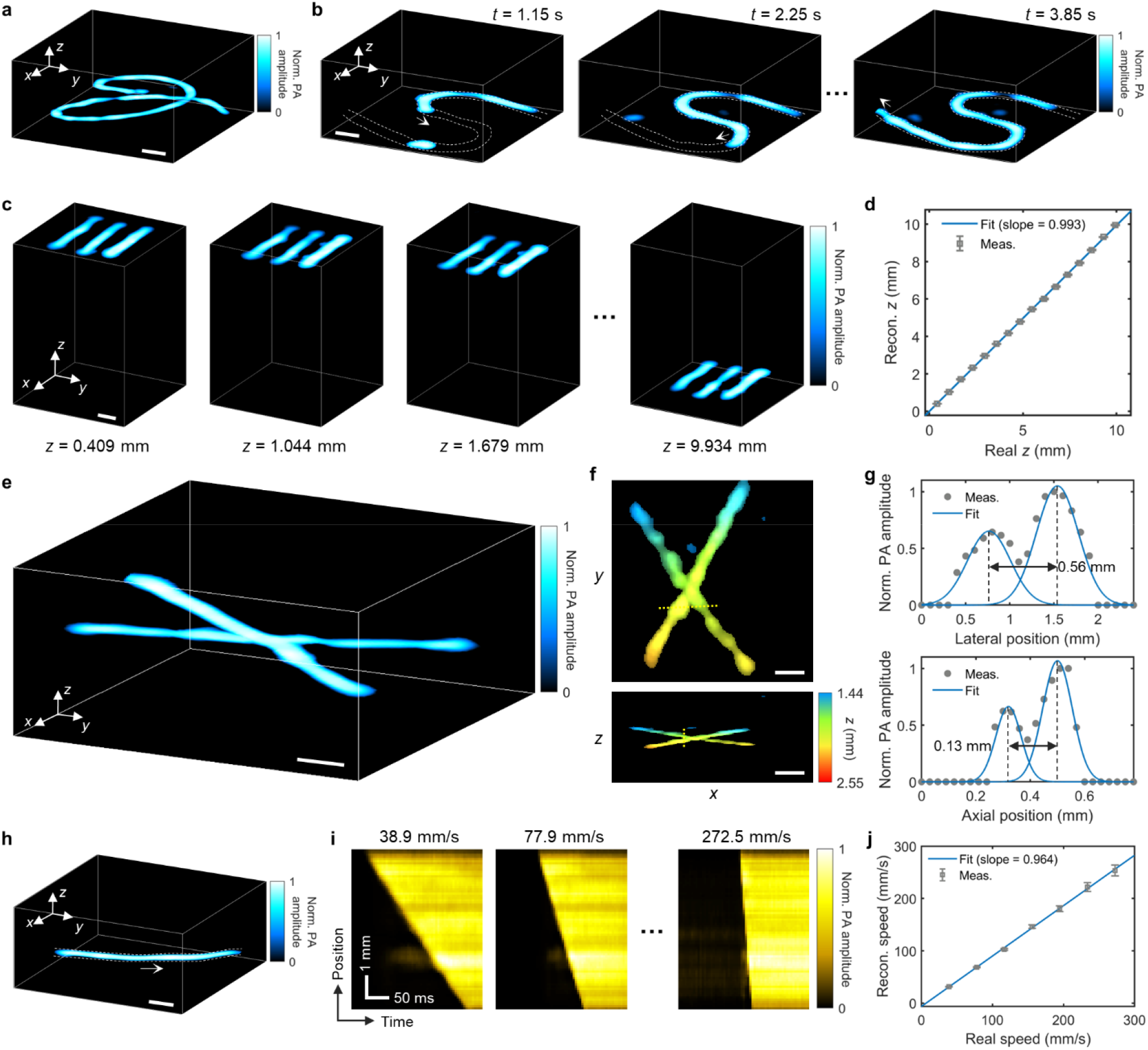
Spatiotemporal characterization of PACTER. **a**, 3D PACTER image of a curved black wire. Norm., normalized. **b**, Snapshots of 4D PACTER showing bovine blood flushing through an S-shaped tube. **c**, 3D PACTER images of three bars printed with black ink on a transparent film. In each image, the object was placed at a different *z* position. **d**, Reconstructed versus real *z* positions of the objects in **c**. The measurement results are plotted as means ± standard errors of the means (*n* = 1,980). The blue curve represents a linear fit. **e**, 3D PACTER image of two crossing human hairs in agarose. **f**, Maximum *z*- (top) and *y*-projections (bottom) of the 3D volume in **e**. The *z* positions of the object are color-encoded. **g**, Profiles along the yellow dashed lines in **f** denoted by gray dots. The blue curves represent two-term Gaussian fits. Black arrows denote the minimum distances that can separate the two hairs. **h**, 3D PACTER image of bovine blood flushing through a tube. The white arrow indicates the flushing direction. **i**, PA amplitudes along the tube in **h** versus time, when the blood flushes through the tube at different speeds. **j**, Speeds of the blood flow quantified from the reconstructed images versus the real speeds in **i**. The measurement results are plotted as means ± standard errors of the means (*n* = 74). The blue curve represents a linear fit. Scale bars, 1 mm.

To quantify the resolution of PACTER, we imaged two human hairs embedded in an agarose block (Fig. 3e). The hairs were intentionally positioned such that they were in close contact with each other, forming a crossing pattern that could be seen in both *z*- and *y*-projections (Fig. 3f). Defining the spatial resolution as the minimum distance that can distinguish the peaks of the two hairs, we found the lateral and axial resolutions of PACTER to be 0.56 mm and 0.13 mm, respectively (Fig. 3g). The anisotropic spatial resolutions along the lateral and axial directions were related to the image formation process in PACTER (Supplementary Notes 4 and 5, Supplementary Figs. 8 and 9). The coarser lateral resolution was due to the acoustic impedance mismatch between the object and the ER (Supplementary Note 4) while both resolutions are limited by the frequency-dependent acoustic attenuation (Supplementary Fig. 10).

To evaluate whether PACTER could be reliably used to image 4D dynamics, i.e., time-lapse movements of 3D objects, we captured 4D images of bovine blood flushing through a tube at different speeds precisely controlled by a syringe pump (Fig. 3h, Supplementary Video 4). Based on the reconstructed 4D images, we plotted the PA amplitudes along the tube (1D images) over time (Fig. 3i), calculated the speeds of the blood flow (Supplementary Fig. 11), and compared them with the real speeds set by the syringe pump (Fig. 3j). A linear relationship (*R^2^* = 0.999) between the reconstructed and real speeds with a slope (0.964) close to 1 can be observed, demonstrating that PACTER is capable of 4D imaging, faithfully reconstructing the dynamics of 3D objects over time. Empowered by the imaging speed of up to a thousand volumes per second, PACTER could resolve the high-speed dynamics of the blood flushing through the tube at 272.5 mm/s in 3D, with a temporal resolution of 1 ms (Supplementary Video 5).

### 4D *in vivo* imaging of mouse hemodynamics with PACTER

Enabled by the capability of noninvasive, label-free, and ultrafast 3D imaging, PACTER is expected to be suitable for monitoring hemodynamics *in vivo*. Here, we evaluated PACTER’s capability in monitoring vital signs in small animals. We imaged the hemodynamics of the abdominal regions of mice (Fig. 4a). With a single laser pulse, PACTER could reconstruct the abdominal vasculature in 3D (Fig. 4b,c). When multiple laser pulses were used, PACTER revealed the 4D dynamics of the blood vessels (Supplementary Videos 6 and 7). Based on the 4D PACTER datasets, we isolated individual blood vessels from the cross sections of the 3D volumes (Fig. 4d,e) and visualized their motions and structural changes (Fig. 4f,g).

**Fig. 4.**
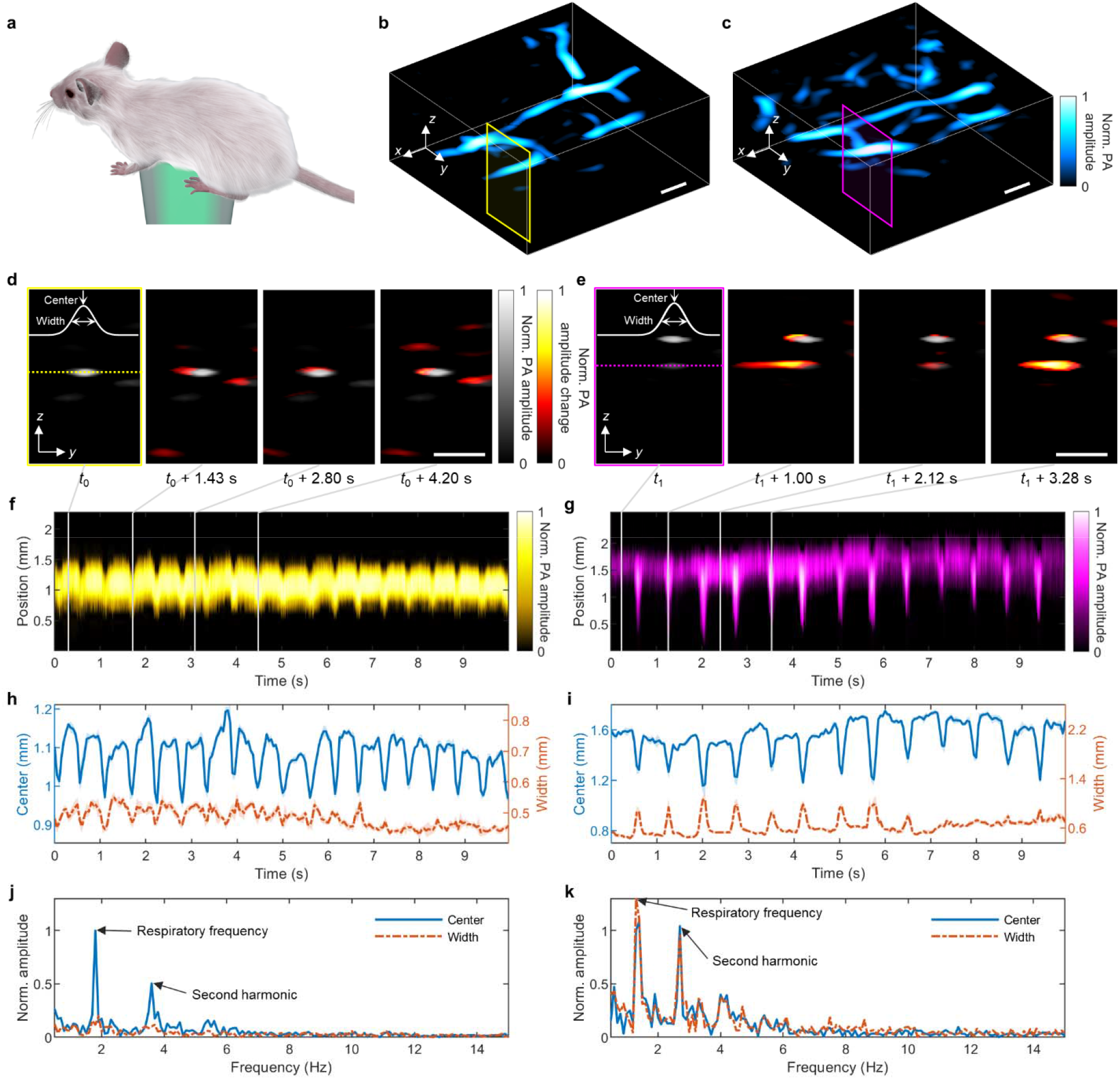
PACTER of mouse hemodynamics *in vivo*. **a**, Schematic of the mouse imaging experiment. **b,c**, 3D PACTER images of the abdominal vasculature of mouse 1 (**b**) and mouse 2 (**c**). Norm., normalized. **d,e**, Cross-sectional 2D images corresponding to the yellow rectangle in **b** (**d**) and the magenta rectangle in **c** (**e**) at four different time instances from the 4D PACTER datasets. *t*_0_ = 0.28 s, *t*_1_ = 0.26 s. White solid curves represent the Gaussian fits of the vessels’ profile denoted by the yellow (**d**) and magenta (**e**) dashed lines. Differences from the first image are highlighted. **f,g**, PA amplitudes along the yellow dashed line (1D images) in **d** (**f**) and the magenta dashed line in **e** (**g**) versus time, where the time instances in **d** and **e** are labeled with vertical gray lines. **h,i**, Center positions (blue solid curves) and widths (orange dash-dotted curves) of the vessels versus time, based on the fits in **d** (**h**) and **e** (**i**). The shaded areas denote the standard deviations (*n* = 5). **j,k**, Fourier transforms of the center positions and widths of the vessels in **h** (**j**) and **i** (**k**), showing the respiratory frequency from the vessel center positions only (**j**) or both the vessel center positions and widths (**k**). Scale bars, 1 mm.

By recording the time-lapse changes of the center positions and widths of the blood vessels, the respiratory motion could be tracked and identified (Fig. 4h,i). Using Fourier analysis, we found that the center position of the blood vessel of mouse 1 fluctuated periodically, exhibiting a respiratory frequency of 1.8 Hz (Fig. 4j), whereas the width of the vessel was relatively stable. In comparison, a respiratory frequency of 1.4 Hz could be observed from both the center position and width of the blood vessel of mouse 2 (Fig. 4k). Further, when we imaged the third mouse (Supplementary Video 8), we observed a respiratory frequency of 1.9 Hz from the width, not the center position, of the blood vessel (Supplementary Fig. 12). The distinct 4D hemodynamics of the blood vessels from the three mice demonstrated that PACTER could be a practical tool in monitoring vital signs, such as breathing, in small animals.

### 4D *in vivo* imaging of hemodynamics in human hands with PACTER

To demonstrate PACTER’s effectiveness in monitoring hemodynamics in humans, we imaged the hand vasculature of two participants. Different regions of the hand, e.g., fingers, proximal phalanx, and thenar regions, were imaged independently as the participants moved their hands to align those regions with the ER (Supplementary Fig. 13). In the following study, we focused on imaging the participants’ thenar vasculature and their responses to cuffing, which was induced by a sphygmomanometer wrapped around the participants’ upper arm (Fig. 5a). Using PACTER, we imaged the thenar vasculature in 3D with single laser pulses (Fig. 5b,c) and reconstructed the 4D dynamics of the blood vessels in response to cuffing (Supplementary Videos 9 and 10). As shown in the maximum amplitude projections of the 4D datasets (Fig. 5d,e), whereas some blood vessels exhibited a relatively stable PA amplitude throughout the experiment, the other vessels showed a decreased PA amplitude after cuffing due to the occlusion of blood flows; when the cuffing was released, the blood flows recovered, and the PA amplitude was rapidly restored (Fig. 5f,g). The different hemodynamics of these two types of blood vessels in response to cuffing may indicate their distinct roles in the circulatory system^47^: the blood vessels with stable and changing PA amplitudes could be venous and arterial, respectively, agreeing with the observations reported in other cuffing-based studies^48–50^. With the capability to simultaneously image both arterial and venous blood *in vivo*, PACTER provides additional benefits over conventional pulse oximetry, which can only monitor arterial blood without spatial resolution^51^.

**Fig. 5.**
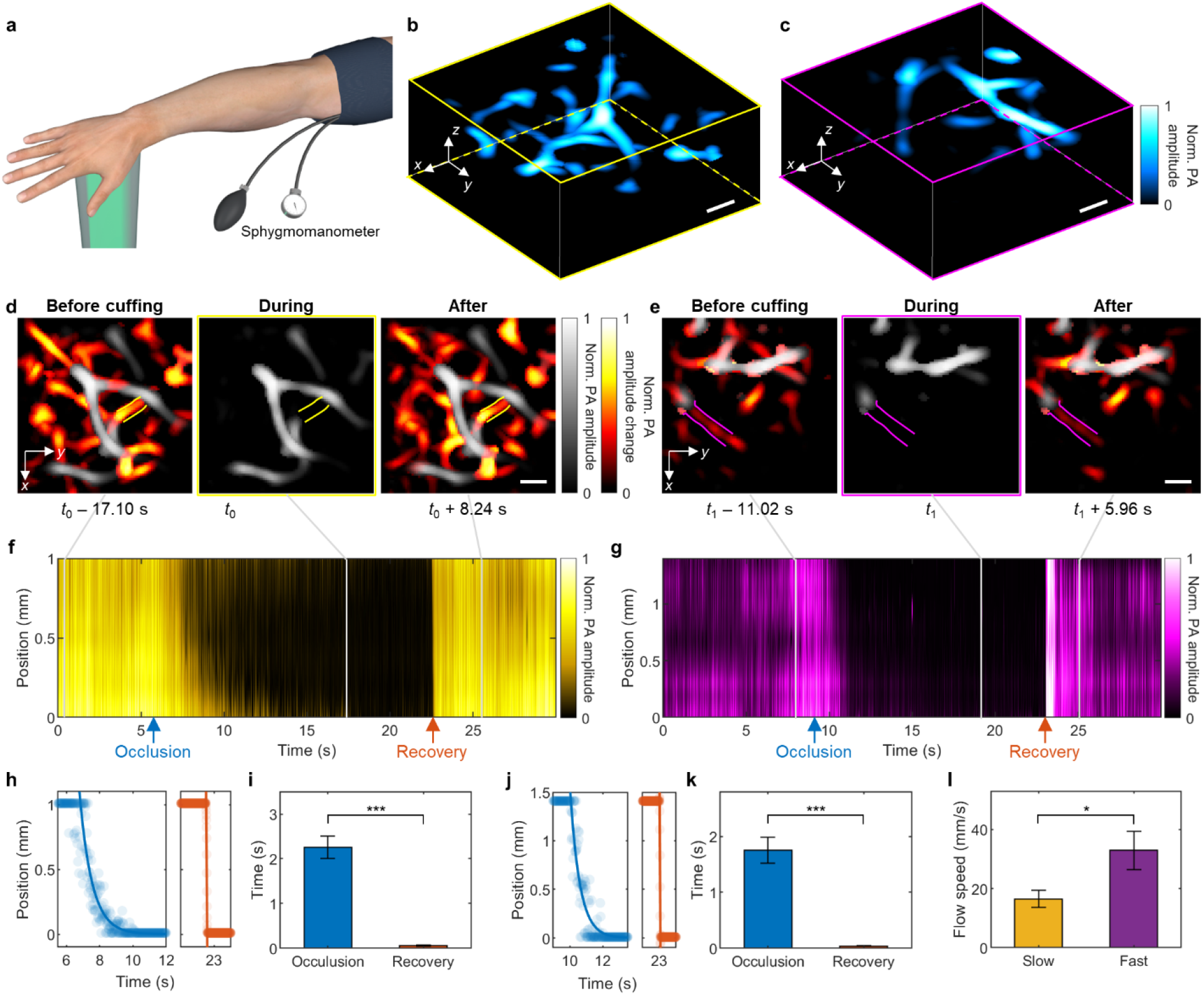
PACTER of human hand hemodynamics *in vivo*. **a**, Schematic of the human hand imaging experiment. **b, c**, 3D PACTER images of the thenar vasculature of participant 1 (**b**) and participant 2 (**c**). Norm., normalized. **d,e**, Maximum amplitude projections of the 3D volumes from the 4D PACTER datasets along the *z* axis in **b** (**d**) and **c** (**e**) at the time instances before, during, and after cuffing. *t*_0_ = 17.44 s, *t*_1_ = 19.02 s. The solid lines flank the vessels under investigation. Differences from the images during cuffing are highlighted. **f,g**, PA amplitudes along the vessels (1D images) in **d** (**f**) and **e** (**g**) versus time, where the time instances in **d** and **e** are labeled with vertical gray lines. The blue and orange arrows indicate peak responses in the occlusion and recovery phases, respectively. **h**, Positions (solid circles) of the blood front along the blood vessel during the occlusion (left) and recovery (right) phases in f. The blue curve is an exponential fit with an occlusion rate of 1.3 ± 0.1 m/s, and the orange curve is a linear fit showing the blood flow speed of 16.5 ± 2.8 m/s. **i**, Comparison between the durations of the occlusion and recovery phases in **f.**\*\*\**P* < 0.001, calculated by the two-sample *t*-test. **j**, Positions (solid circles) of the blood front along the blood vessel during the occlusion (left) and recovery (right) phases in **g**. The blue curve is an exponential fit with an occlusion rate of 2.4 ± 0.3 m/s, and the orange curve is a linear fit showing the blood flow speed of 32.9 ± 6.5 m/s. **k**, Comparison between the durations of the occlusion and recovery phases in **g**. \*\*\**P* < 0.001, calculated by the two-sample *t*-test. **l**, Comparison between the blood flow speeds during recovery in **f** and **g**. \**P* < 0.05. Scale bars, 1 mm.

Because PA amplitudes have 100% sensitivity to optical absorption^11^, the 4D hemodynamics imaged by PACTER revealed the real-time changes in the blood vessels in response to cuffing, and the linear position of the blood front during the recovery phase could be used to measure the blood flow speed^50^. For participant 1, the occlusion rate of the vessel was found to be 1.3 ± 0.1 m/s, significantly slower than the blood flow speed of 16.5 ± 2.8 m/s extracted from the recovery phase (Fig. 5h,i). For participant 2, the occlusion rate and the blood flow speed of the vessel were found to be 2.4 ± 0.3 m/s and 32.9 ± 6.5 m/s, respectively (Fig. 5j,k), exhibiting a greater blood flow speed compared with participant 1 (Fig. 5l). Immediately after an imaging session, we asked participant 1 to slightly move their hand and used PACTER to image a different area of the thenar region (Supplementary Video 11). Using the same analysis on a different blood vessel, the occlusion rate and the blood flow speed were found to be 0.6 ± 0.1 m/s and 22.4 ± 6.4 m/s, respectively (Supplementary Fig. 14). Taken together, we demonstrated that PACTER could monitor the hemodynamics in human, including the consistent responses of thenar vasculature to cuffing, and capture the variability in blood flow speeds.

## Discussion

Although the current implementation of PACTER requires motorized stages for calibration, a pulsed laser for illumination, and a DAQ card for data acquisition, these requirements could be fulfilled using cheaper and more compact alternatives. Owing to the universal calibration capability, the ER in PACTER could be pre-calibrated, eliminating the need for motor-based calibration. The pulsed laser and the DAQ card could be replaced with cost-effective light-emitting diodes (LEDs)^52^ and microcontrollers^53^, respectively, which could further enhance the portability of the system. Additionally, mass production of the ER and the single-element ultrasonic transducer could substantially lower the cost of the system, making PACTER more accessible to users or researchers in low-resource settings, further lowering the barriers to clinical translation.

Due to the large dimensions of the ER compared with the acoustic wavelength, the PA waves need to propagate a long distance inside the ER; therefore, a slight change in the speed of sound due to temperature fluctuations^54^ would cause large differences in the measured PACTER signal (Supplementary Video 12). To address this problem, we built a temperature stabilizing box to maintain the temperature of the ER (Supplementary Fig. 15), which stabilizes the temperature of the ER at a set temperature, e.g., 30 °C, at all times, guaranteeing a constant speed of sound throughout the experiments. In addition, the penetration depth of our PACTER system was limited to 3.6 mm *in vivo* due to the strong attenuation of 532 nm light by endogenous chromophores in biological tissue^55^. Changing the wavelength to 1064 nm could increase the penetration depth to several centimeters^15,20^. Another limitation of PACTER is the relatively small FOV (8 mm x 8 mm) constrained by the diameter of the glass rod in the ER. We believe that the design of the ER could be optimized further to achieve a larger FOV, enabling new applications such as vascular biometrics^56^. Finally, the spatial resolution of PACTER is currently limited by the acoustic impedance mismatch between the object and the ER, which could be addressed in the future by adding an impedance matching layer on top of the ER.

In summary, PACTER, a noninvasive, label-free, and ultrafast imaging technique, enables 4D imaging of hemodynamics in humans using the 1D signal captured by a single detector, achieving an imaging speed of up to a thousand volumes per second. We have demonstrated PACTER’s capability to visualize the 4D hemodynamics in humans and small animals. We have also shown the convenience of using PACTER to image different objects, including human hands and mouse abdomens, without the need for recalibration. PACTER’s high imaging speed allows for immediate intervention in case of abnormal hemodynamic changes. Additionally, PACTER’s low cost and compact form factor are ideal for point-of-care testing, facilitating quick and easy assessment of hemodynamic parameters at the bedside or in remote locations. We envision that PACTER will have profound impacts on a wide range of applications in biomedical research and clinical settings, including home care of diabetic-foot ulcers^37^ or carotid-artery disease^38^, point-of-care screening for hypertension^39^, and simultaneous oximetry of both arterial and venous blood in intensive care units^40^. Furthermore, PACTER’s single-shot 3D imaging concept using a single-element detector can extend beyond optical imaging, aiding fields such as medical ultrasonography^41^, underwater sonar^42^, and airborne radar^43^.

## Methods

### Experimental setup

In the PACTER system, the power of a 5-ns pulsed laser beam at 532 nm (INNOSLAB IS8II-DE, EdgeWave; 1 kHz pulse repetition rate) was controlled by a half-wave plate (WPH10M-532, Thorlabs) and a polarizing beam splitter (PBS25-532-HP, Thorlabs). The beam reflected by the PBS was sent to a beam trap (BT610, Thorlabs). The beam transmitted through the PBS was expanded by a beam expander consisting of two lenses (ACN254-050-A and AC254-100-A, Thorlabs). During calibration (Fig. 1a), the expanded beam was steered by the mirrors mounted on two motorized linear translation stages (PLS-85, PI), which were controlled by two motor drivers (CW215, Circuit Specialists). The beam diameter was adjusted to be 2 mm by an iris (SM1D12, Thorlabs), and the beam was sent through a lens (AC254-300-A, Thorlabs) and focused on top of the ER. A container with a window at its bottom sealed with an optically and ultrasonically transparent disposable polyethylene membrane was filled with bovine blood, which was used as a uniform optical absorber for calibration (Fig. 2a). The container was placed on top of the ER, where ultrasound gel (Aquasonic 100, ParkerLabs) was applied between the polyethylene membrane and the ER surface to facilitate acoustic coupling. During imaging (Fig. 1b), the beam diameter was adjusted to be 6 mm by the iris, and the beam was sent through a fly’s eye homogenizer (Supplementary Note 1) consisting of two microlens arrays (#64-480, Edmund Optics) and a lens (AC254-250-A, Thorlabs), which homogenized the beam in the imaging volume. Ultrasound gel was applied between the object and the top of the ER to facilitate acoustic coupling. The PACTER signals detected by the ultrasonic transducer were amplified by two low-noise amplifiers (ZKL-1R5+, Mini-Circuits), filtered by a low-pass filter (BLP-70+, Mini-Circuits), and digitized by a data acquisition card (ATS9350, AlazarTech) installed on a desktop computer. A multifunctional input/output (I/O) device (PCIe-6321, National Instruments) was used to control the laser, the motorized stages, and the data acquisition card.

### Universally calibratable ER

The ER in PACTER consists of a right-angle prism (PS611, Thorlabs; 25 mm right-angle edge length) and a customized optical rod (VY Optoelectronics; 18 mm diameter, 175 mm length, top and bottom surfaces polished to 60-40 surface quality). Both components were made of ultraviolet (UV) fused silica, which has good optical transparency and low acoustic attenuation. To enhance ergodicity^57^, the edges of the prism were ground by a diamond saw (SYJ-150, MTI Co.) following a sawtooth pattern to obtain chaotic boundaries^36^. The prism and the rod were glued by UV-curing optical adhesive (NOA68, Norland Products), following exposure under UV light for 12 hr. To avoid the change of speed of sound due to temperature fluctuations during experiments, the whole ER was sealed in a temperature stabilizing box (Supplementary Fig. 15) regulated by a thermocouple (SA1-E, Omega Engineering), a heating pad (SRFG-303/10, Omega Engineering), and a temperature controller (Dwyer 32B-33, Cole-Parmer).

### Fabrication of the ultrasonic transducer

The fabrication process of the ultrasonic transducer is as follows. First, PMN-PT piezoelectric single crystal (CTS Corporation) was chosen as the core component for acoustic-electrical conversion due to the excellent piezoelectric coefficient and high permittivity, which is suitable for high-frequency transducers with small aperture sizes because of the general electrical impedance matching (50 ohms). Second, based on the material parameters, a transducer modeling software (PiezoCAD) based on Krimboltz, Leedom, and Mattaei (KLM) equivalent circuit model was employed to simulate and optimize the design of the transducer. Hence, a 40 MHz PMN-PT transducer with a small active aperture size of 0.4 × 0.4 mm^2^ was designed and obtained. The piezoelectric element exhibits a central frequency of 40 MHz. Third, the piezoelectric crystal was lapped to the required thickness (40-μm), gold electrodes were sputtered on both sides, and then a layer of conductive silver paste (E-solder 3022) was deposited onto the piezoelectric sheet as a backing layer. Fourth, the acoustic stack was diced into designed elements size. Fifth, using Kapton tapes as a mask, a gold electrode was sputtered on the corner of the hypotenuse surface of the prism of the ER (Supplementary Fig. 3). The piezoelectric element was then affixed directly to the electrode on the ER by using a thin layer of conductive silver paste. The wires were connected out to read the signals. Last, a thin parylene layer as the protective layer was deposited onto the device.

### Data acquisition

A custom-written LabVIEW (National Instruments) program was used to trigger the pulsed laser, drive the motorized stages during calibration, and acquire the data. The PACTER signals were acquired at a sampling rate of 250 megasamples per second, and a sampling length of 65,532 data points per acquisition. Due to the distance between the object and the ultrasonic transducer, a 28-μs delay was added to the data acquisition following the laser trigger. During calibration, to improve the signal-to-noise ratio (SNR) of the signal, we repeated the acquisition 500 times at each calibrated virtual transducer and used the averaged signal for PACTER reconstruction. To prevent motor backlash, the data were acquired only when the motor was moving forward; the acquisition stopped when the motor returned. During imaging, to improve the temporal resolution of the system, no signal averaging was used, and motor scanning was disabled.

### Imaging protocols

All human and animal imaging experiments were performed in accordance with relevant guidelines and regulations. The human imaging experiments followed the protocol approved by the Institutional Review Board (IRB) of the California Institute of Technology. Two healthy adult participants were recruited for this study. Written informed consent was obtained from both participants according to the protocol. The PACTER system was pre-calibrated using bovine blood, and the participants were enrolled for the imaging procedure only. For imaging the vasculature in human hands, after applying ultrasound gel, the participants were instructed to place their hands on top of the ER. For imaging the human hand hemodynamics in response to cuffing, a sphygmomanometer was wrapped around the participants’ upper arm (Fig. 5a). To induce blood vessel occlusion, the sphygmomanometer was inflated to high pressure (200 mgHg), maintained for a short time (~15 s), and then quickly released; the total imaging time was 30 s. The animal imaging experiments followed the protocol approved by the Institutional Animal Care and Use Committee (IACUC) of the California Institute of Technology. IRB and IACUC were aware of both protocols before approval. The fluence of the laser beam for imaging (5 mJ/cm^2^) was within the American National Standards Institutes (ANSI) safety limit for laser exposure (20 mJ/cm^2^ at 532 nm)^58^.

### Animal preparation

Female athymic nude mice (Hsd: Athymic Nude-Foxn1nu, Envigo; 15–20 g and 4–5 weeks old) were used for the animal imaging experiments. Before imaging, the mouse was placed in a small chamber with 5% vaporized isoflurane mixed with air for anesthesia induction. It was then transferred to a customized animal mount, which has a hole at the bottom such that the abdomen of the mouse can be imaged by PACTER. Ultrasound gel was applied on top of the ER, and then the animal mount was lowered until the mouse abdomen was in contact with the ER. Throughout the imaging session, the mouse was kept anesthetized with a continuous supply of 1.5% vaporized isoflurane, its head was fixed to the stereotaxic frame of the mount, and its body temperature was maintained at ~38 °C by a heating pad.

### Sample preparation

For calibration, refrigerated bovine blood (#910, Quad Five) was restored to room temperature and transferred to the customized container (Fig. 2a) for acquiring the calibration signals for PACTER reconstruction. Besides bovine blood, black tape (6132-BA-10, 3M), black rubber (5508T44, McMaster-Carr), and black ink (X-1, Tamiya) were also used to test their performance as uniform optical absorbers for calibration. During the test, black tape and black rubber were cut into 10 mm x 10 mm sheets and placed on top of the ER, with ultrasound gel as the coupling medium; black ink was stored in the customized container. For imaging, black wires (8251T9, McMaster-Carr) were bent into curved shapes and placed in the customized container filled with water. Bar patterns were printed with black ink on a transparent film (ITF-30, Octago), cut into 10 mm x 10 mm sheets, and placed in the water-filled customized container; a wooden stick mounted on a manual linear translation stage (PT1, Thorlabs) was glued to the back of the film to control its *z* position in the imaging volume. Human hairs were embedded in a 4% agarose block (A-204-25, GoldBio), and the block was placed in the water-filled customized container during imaging. Polyurethane tubes (MRE025, Braintree Scientific; 0.012” inside diameter) were first placed in the water-filled customized container in straight or curved shapes, and a syringe was used to flush bovine blood through the tubes; the speed of the blood flow was controlled by a syringe pump (NE-300, New Era).

### PACTER reconstruction

In PACTER, the signal *s*(*t*) detected by the ultrasonic transducer at time *t* in a homogeneous medium is expressed as (Supplementary Note 3)

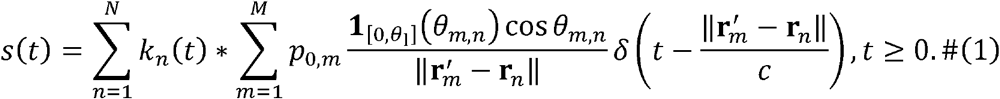

Here, *M* and *N* are the numbers of the source points and the calibrated virtual transducers, respectively; *k_n_* (*t*) is the normalized impulse response from the calibration at the *n*-the virtual transducer; 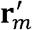 and *r_n_* are the locations of the *m*-th source point and the *n*-th virtual transducer, respectively; *p_0,m_* is a value proportional to the initial pressure at 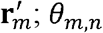 denotes the incidence angle satisfying 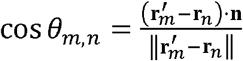 with **n** being the normal vector of the calibration plane; θ_1_ is the critical angle the ultrasonic refraction from water to fused silica; **1**_[0,*θ*_1_]_ represents the indicator function defined in Eq. (S6); *c* is the speed of sound in the homogeneous medium; *δ*(*t*) denotes the delta function.

Discretizing Eq. (1), we obtain the forward model

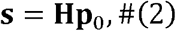

where **s** represents a vector of length *L*, **p**_0_ denotes a vector of length *M*(=*M*_1_*M*_2_*M*_3_) which consists of all voxels in a 3D image of size *M*_1_ × *M*_2_ × *M*_3_, and **H** is the system matrix of size *L* x *M*. This forward model has a computational complexity of max{*O*(*MN*), *O*(*NL*log_2_*L*)}. To obtain an image from the signals **s**, we invert the forward model by solving the regularized optimization problem

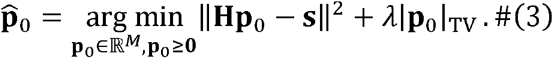

Here, |**p**_0_|_TV_ denotes the total variation of the 3D image corresponding to **p**_0_, and *λ* is the regularization parameter. Numerically, we solve this optimization problem through a Fast Iterative Shrinkage-Thresholding Algorithm (FISTA)^59^.

### Image processing

The reconstructed images were first denoised using a 3D median filter (in the 3-by-3-by-3 neighborhood) and smoothed using a 3D Gaussian filter (with a 0.1-by-0.1-by-2 standard deviation kernel). We then applied a Hessian-matrix-based vesselness filter^60^ to the denoised images to improve the contrast of vascular structures in 3D. Finally, we added the vesselness-enhanced images (self-normalized) with a weighting factor of 0.8 back to the filtered images with a weighting factor of 0.2 and obtained the presented images. The images were rendered in 3D or 4D (time-lapse 3D) using the Imaris (Bitplane) software. The speeds of bovine blood flushing through the tube in Fig. 3 were calculated by differentiating the PA amplitudes along the tube and fitting the relationship between the traveling distance of the blood front versus time. The vessels’ profiles in Fig. 4 were fitted as a Gaussian function, exp(–(*x* – *x*_0_)^2^/*w*^2^), where the center positions and widths of the vessels were estimated from *x*_0_ and 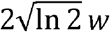, respectively. The vessels’ profiles in Supplementary Fig. 12 were fitted as a two-term Gaussian function, where the center positions and widths were estimated from the first term. The positions of the blood front along the blood vessels in Fig. 5 and Supplementary Fig. 14 were obtained through thresholding the PA amplitude profiles. Denoting the total length of the blood vessel profile as *L_p_*, we fitted the positions of the blood front along the blood vessels during the occlusion and recovery phases with *d*_o_(*t*) = *a* exp(–*v_o_t*/*L_p_*) and *d_r_*(*t*) = –*v_r_t* + *b*, respectively, where *a* and *b* were constants, *v_o_* was the occlusion rate, and *v_r_* was the blood flow speed. The durations of the occlusion and recovery phases were estimated by *t_o_* – 3*L_p_*/*v_o_* and *t_r_* = 0.93*L_p_*/*v_r_*, respectively, which were the time it took for the blood front to propagate 95%, i.e., 1 – exp(–3), of *L_p_*. To determine if the differences between *t_o_* and *t_r_* were significant, we applied a Welch’s (unequal variances) *t*-test to determine the *P* values under the null hypothesis that the mean values of *t_o_* are not different from those of *t_r_*. The same *t*-test was also performed to determine if the difference between the blood flow speeds in Fig. 5l was significant.

## Data availability

The main data supporting the results in this study are available within the paper and its Supplementary Information. Other data are too large to be publicly shared, yet they are available for research purposes from the corresponding author on reasonable request.

## Code availability

The reconstruction code, the system control software, and the data collection software are proprietary and used in licensed technologies, yet they are available from the corresponding author upon reasonable request.

## Acknowledgments

We thank Yanyu Zhao for contributing to the universal calibration. This work was supported in part by National Institutes of Health grants R01 EB028277, U01 EB029823, and R35 CA220436 (Outstanding Investigator Award). The computations presented here were conducted in the Resnick High Performance Computing Center, a facility supported by Resnick Sustainability Institute at the California Institute of Technology

## Author contributions

Y.Zhang and L.V.W. conceived and designed the study. Y.Zhang, L.L., R.C., and K.M. built the imaging system. Y.Zhang developed the data acquisition program. P.H. developed the 3D reconstruction algorithm. Y.Zhang, L.L., R.C., and A.K. performed the experiments. Y.Zhang, P.H., and X.T. processed and analyzed the data. Y.Zeng, L.J., and Q.Z. fabricated the ultrasonic transducer. L.V.W. supervised the study. All authors contributed to the writing of the manuscript.

## Supplementary Information

### Supplementary Note 1 Fly’s eye homogenizer in PACTER

We use a fly’s eye homogenizer to provide uniform illumination for PACTER^61^. As shown in Supplementary Fig. 2, the two microlens arrays form multiple parallel Köhler illumination systems side-by-side. The original beam entering the first microlens array is divided into multiple beamlets. Through the lenslets pairs in the microlens arrays and the spherical lens, each beamlet of the original beam is imaged to the homogenization plane, i.e., the top of the ER. Because the images of the beamlets are all superimposed on the homogenization plane, the intensity differences among the beamlets disappear in their superimposed images. Therefore, the intensity distribution of the homogenized beam is independent of the homogeneity of the original beam. Further, the square-type microlens arrays in the setup will generate a square flat-top intensity distribution in the homogenization plane, which provides a good match for the square-type calibration pattern in PACTER, allowing accurate mapping between the calibration and imaging areas.

The width of the homogenized beam, *d_H_*, is given by

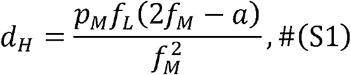

where *p_M_* and *f_M_* are the pitch and focal length of the lenslets in the two identical microlens arrays, *f_L_* is the focal length of the spherical lens, and *a* is the separation between the microlens arrays. In PACTER, *a* is set to be identical to *f_M_*, leading to

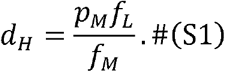

The divergence half-angle after the homogenization plane, *θ*, is given by

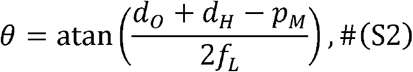

where *d_0_* is the diameter of the original beam. In PACTER, we use the microlens arrays with *p_M_* = 0.5 mm and *f_M_* = 15 mm, the spherical lens with *f_L_* = 250 mm, and the original beam with a diameter of *d*_0_ = 6 mm. Therefore, the homogenized beam has a width of *d_H_* ≈ 8 mm, matching the size of the calibration pattern (80 by 80 steps with a step size of 0.1 mm), and the divergence half-angle *θ* ≈ 2°. The small divergence ensures homogenous illumination across the whole 3D volume (8 mm x 8 mm x 3.6 mm) for *in vivo* imaging. Within the 3.6 mm depth, the illumination beam merely diverges laterally by 0.13 mm, which is much smaller than the lateral resolution (0.56 mm) of PACTER. Hence, the beam divergence within the imaging volume can be ignored.

### Supplementary Note 2 Universal calibration of PACTER

Recently, we developed photoacoustic topography through an ergodic relay (PATER), which captured a wide-field PA image with a single laser shot using a single-element ultrasonic transducer^34–36^. However, PATER suffered from the following limitations. (1) PATER could only image a 2D projection of the object, unable to capture a tomographic image in 3D. (2) The ER in PATER had to be re-calibrated for each different object; because the calibration relied on a point-by-point scanning of a laser beam, the imaging procedure was time-consuming. (3) PATER required that the boundary conditions of the object and the ER stay unchanged throughout the experiment and thus could not be used for long-term imaging in an unstable environment.

One major distinction between PATER and PACTER lies in the structures of their ERs. In PATER, the ER is simply a prism (Supplementary Fig. 5a). After a PA signal is generated from the object (*t*_0_) and detected by the transducer (*t*1, it quickly propagates to and reflects from the boundary between the object and the ER (*t*_2_), as shown in the simulation based on the *k*-wave toolbox^62^ (Supplementary Fig. 5b). Once the boundary condition between the object and the ER changes because of movements of the object, instability of the system, or switching to a different object, the signal will also change. Therefore, if the ER is calibrated with one object, e.g., bovine blood, and then used to image a different object, e.g., a black wire, the reconstruction will fail (Supplementary Fig. 5c). In other words, the PATER signal is strongly object-dependent (Supplementary Fig. 5d), and it requires that the boundary conditions of the object and the ER remain unchanged throughout the experiment.

In comparison, the ER of PACTER consists of a prism and a fused silica rod, where the rod functions as an acoustic delay line that temporally separates the initial and reflected PA signals (Supplementary Fig. 5e). When a PA signal is generated from the object (*t*_0_) and detected by the ultrasonic transducer (*t*_1_, a part of the acoustic signal is trapped and scrambled in the prism, whereas the other part is reflected toward the object and then reverberated to the transducer (*t*_2_), as denoted by the black dotted arrows in the simulation (Supplementary Fig. 5f). Because only the reflected part of the acoustic signal will be affected by the boundary condition between the object and the ER, once this part is excluded from the measurement, the acquired signal will be object-independent. Consequently, the ER can be used to image any object, e.g., a black wire, despite it has been calibrated only once using a different object, e.g., bovine blood (Supplementary Fig. 5g). Whereas almost the whole PATER signal is object-dependent (Supplementary Fig. 5d), a large segment (>100 μs) of the PACTER signal is object-independent (Supplementary Fig. 5f), enabling universal calibration in PACTER despite its 3D imaging capability (Supplementary Video 2).

### Supplementary Note 3 Forward model and image reconstruction

We performed calibrations at pixels on a 2D plane and used these pixels as virtual ultrasonic transducers for 3D imaging. If non-zero initial pressure exists only on the calibration plane, the detected signal *s*(*t*) at time *t* can be expressed as

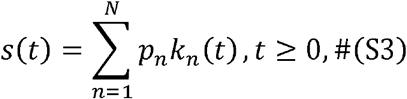

where *N* is the number of calibrated virtual transducers, *k_n_*(*t*) is the normalized impulse response from the calibration at the *n*-th virtual transducer, and *p_n_* is the root-mean-squared PA amplitude proportional to the initial pressure at the *n*-th virtual transducer.

For initial pressure in a 3D volume, we assume *M* source points located at 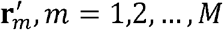, in an acoustically homogeneous 3D region attached to the calibration plane. The PA wave generated from the source point at 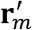 propagates to the calibrated virtual transducer ***r**_n_* with the speed of sound *c* after time 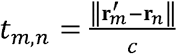, which, through the ER, adds 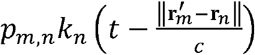 to the detected signal, with the PA amplitude *p_m,n_* quantified as 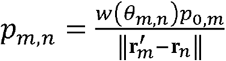. Here, *θ_m,n_* denotes the incidence angle satisfying cos 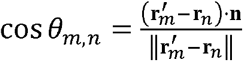 with **n** being the normal vector of the calibration plane; function *w*(*θ_m,n_*) describes a virtual transducer’s angle-dependent sensitivity; and *p_0,m_* is proportional to the initial pressure at 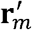. We replace *p_n_k_n_*(*t*) in Eq. (S3) with 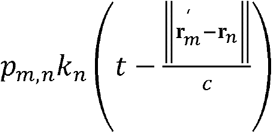 from all the *M* source points and obtain the detected wide-field PA signal

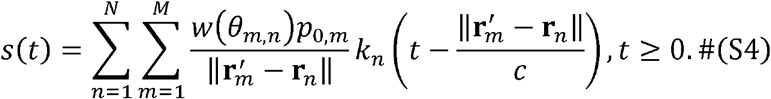

Here, we define *k_n_*(*t*) = 0, *n* = 1,2…, *N*, *t* < 0. For sufficiently small virtual ultrasonic transducers, we assume that

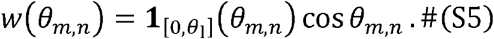

Here, we use the indicator function

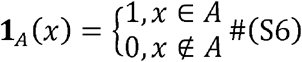

to rejection detections with incidence angles greater than the critical angle *θ*_1_ (quantified in Supplementary Note 4). Substituting Eq. (S5) into Eq. (S4) yields

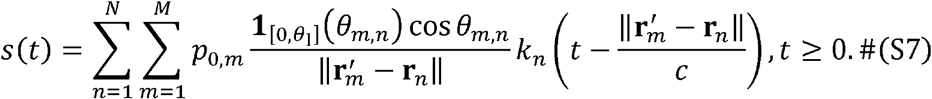

We let *L* be the number of time points after temporal discretization. Then the computational complexity of a forward model based on Eq. (S7) is *O*(*MNL*).

To accelerate the forward model in Eq. (S7), we split the delay term 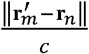 from function *k_n_*(*t*) through temporal convolution:

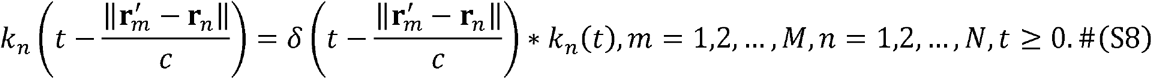

Substituting Eq. (S8) into Eq. (S7), we obtain

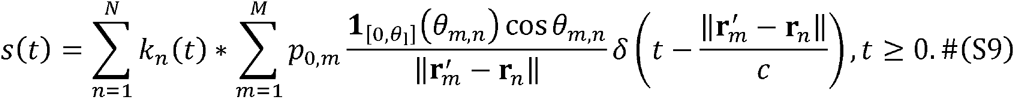

The inner summation in Eq. (S9) has a complexity of *O*(*MN*) and each temporal convolution is implemented through three fast Fourier transforms (FFTs) with a complexity of *O*(*L*log_2_*L*). Thus, the forward model based on Eq. (S9) has a computational complexity of max{*O*(*MN*), *O*(*NL*log_2_*L*)}, which is 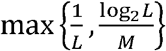 times of *O*(*MNL*). In this work, *L* = 65,536, *M* = 80 × 80 × 120, and 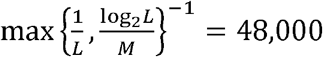, which indicates the factor of acceleration.

### Supplementary Note 4 Spatial resolution analysis based on the virtual transducer array

We analyze the anisotropy of the spatial resolution from two perspectives: one with an emphasis on the physical intuition (this Note), the other with an emphasis on the mathematical derivation (Supplementary Note 5). The two perspectives clarify the differences in value between the lateral and axial resolutions.

We first analyze the anisotropy of the spatial resolution by assuming that the PACTER signals detected by the single-element transducer are accurately decoded to the signals detected by the 80 × 80 virtual transducers. We use the spectrum of the PACTER signal in Supplementary Fig. 8a, denoted as *ŝ*(*f*), to approximate the detection spectrum of each virtual transducer element.

From *ŝ*(*f*), we obtain the central frequency 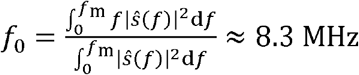 MHz and the bandwidth 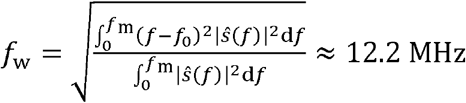 MHz, respectively^63^. Here, we set the upper frequency as *f*_m_ = 125 MHz, which is the Nyquist frequency corresponding to the 250 MHz sampling rate. Additionally, the speed of sound in fused silica (longitudinal wave speed *c*_1_ = 5.9 mm · μs^-1^)^64^ is greater than that in water (*c* = 15 mm ·μs^-1^)^65^, leading to a critical angle (denoted at *θ*_1_) of the ultrasonic wave’s refraction from water into fused silica (Supplementary Fig. 8b). The critical angle satisfies 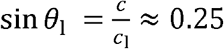, which equals the numerical aperture (NA) of the virtual 2D array. Here, the longitudinal wave speed is used to quantify the critical angle due to the single-element transducer’s dominant sensitivity to the longitudinal waves^66^. For the virtual 2D array, the axial resolution (along the *z*-axis) is quantified as 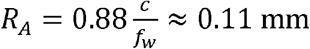, while the lateral resolution (in the *x-y* plane) is quantified as 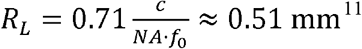. Both values are close to the measured axial (0.13 mm) and lateral (0.56 mm) resolutions shown in Fig. 3g.

### Supplementary Note 5 Spatial resolution analysis based on the system matrix

Obtaining signals detected by all the virtual transducers is an ill-posed problem and we reconstruct an image directly from the single-element-detected signals. We iteratively invert the system matrix **H** (of size *L* X *M*) to obtain the images, which manifest anisotropic resolutions (see Methods). To explain the anisotropy in theory, we analyze the gradient matrix **H^T^H**, whose property eventually determines the image resolution in the iterative method. The *m*-th column of **H** is a temporally discretized form of the following transducer’s response to the point source at
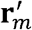:

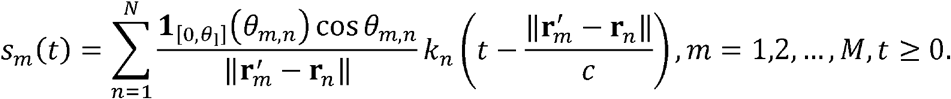

Accordingly, an element at the *m*_1_-th row and *m*_2_-th column in **H^T^H** is expressed in temporally continuous form as

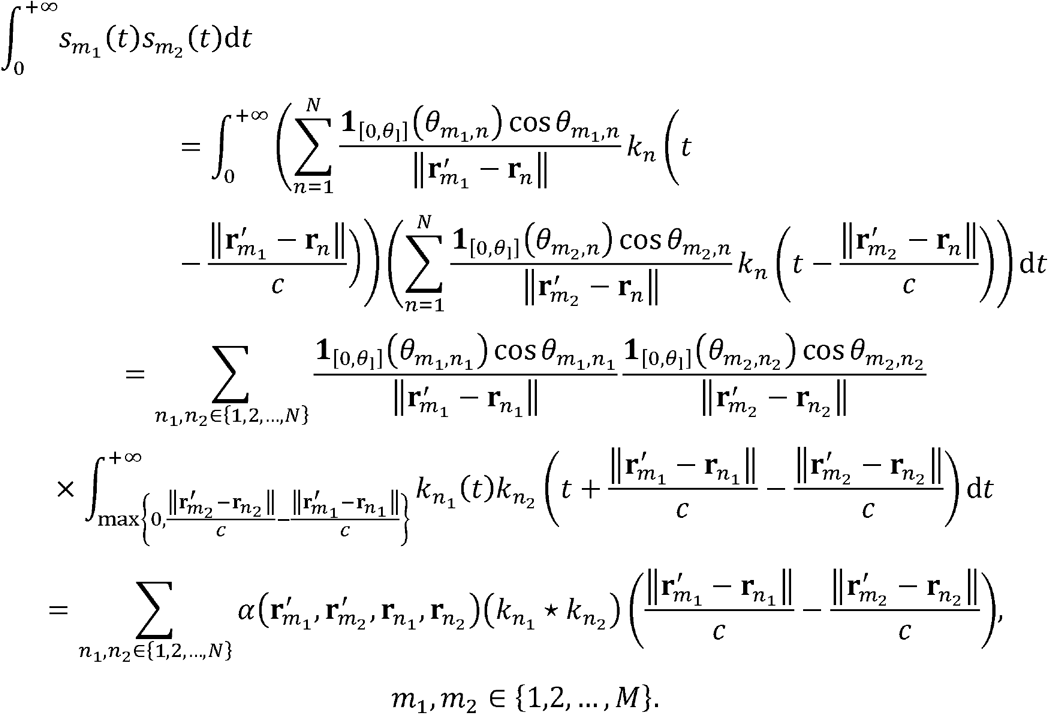

Here, we define a weighting factor

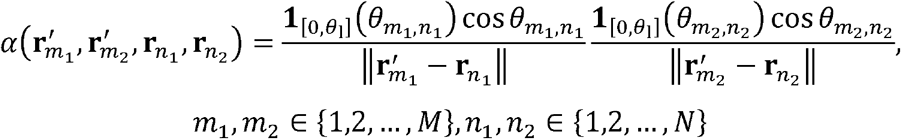

and a cross-correlation function

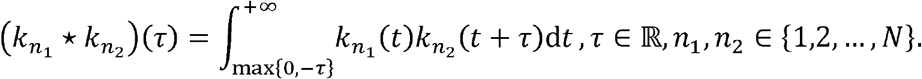

For any non-zero response *s_m_*(*t*), we have 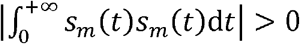. A practical imaging system requires 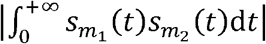 to be small for any *m*_1_ ≠ *m*_2_. The image resolution is determined by the decay speed of 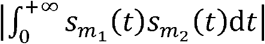 as (*m*_1_, *m*_2_) moves away from the diagonal of **H^T^H**.

To explain the anisotropy of the resolution at 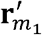, we fix *m*_1_ and vary *m*_2_ so that 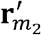 moves from 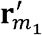 to other locations around 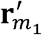. Meanwhile, we estimate the decay of 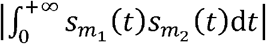. We confine the movement of 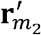 in a small region around 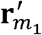 such that the change of 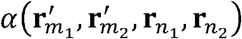 is negligible. Thus, we only need to elaborate on the decay of 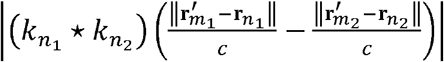.

We first discuss two special cases: the maximum-cross-correlation matrix

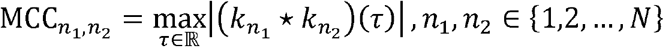

and the autocorrelation function (*k_n_* ✶ *k_n_*)(*τ*), *n* = 1,2,…, *N*. Both functions are determined by the bandwidth of the transducer and the configuration of the ER. Here, we performed calibrations at *N* = 80 × 80 pixels with a spacing of 0.1 mm in each dimension. We pick two locations 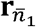 and 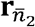 and calculate the values of 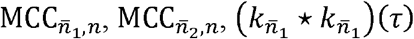, and 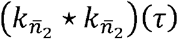. The values of 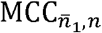 and 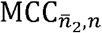 for different *n* form two images shown in Supplementary Fig. 9a and b, respectively. In each image, we draw two perpendicular lines *L*_1_ and *L*_2_ centered at the corresponding calibration location and compare the pixel values along the lines with the autocorrelation function values (*AC*) expressed as 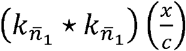 or 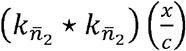, as shown in Supplementary Fig. 9c and d, respectively. As a generalization of the full width at half maximum (FWHM), we define *FW_F,β_* as the full width at *β* times the maximum of a line profile *F*, and 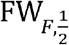 is equivalent to the FWHM of *F*. For the first selected calibration location 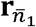, the decay of 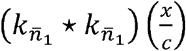 vs. *x* and the decay of 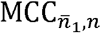 vs. 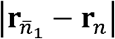 are shown in the plots *AC* and *L*_1_-*L*_2_, respectively, in Supplementary Fig. 9c. We calculated *FW_AC,β_*, *FW*_*L*_1_,*β*_, and *FW*_*L*_2_,*β*_ for *β* = 0.20 and 0.40. The values are shown in Supplementary Fig. 9c. We repeat the quantification for the second selected location 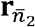, as shown in Supplementary Fig. 9d. We observe that the increase of either 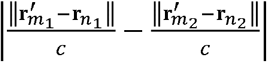 or |**r**_*n*_1__ – **r**_*n*_2__| causes 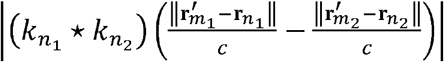 to decay. Equivalently, avoiding the decay of 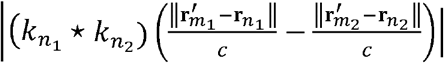 requires both 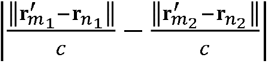 and |**r**_*n*_1__ – **r**_*n*_2__| to be small.

Next, we analyze how the probability of both 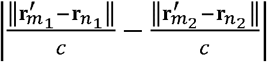 and |**r**_*n*_1__ – **r**_*n*_2__ | being small decreases as 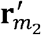 moves away from 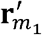. We only discuss two representative cases: (1) 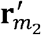 moves on the plane crossing 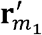 and parallel to the calibration plane (Supplementary Fig. 9e and f); (2) 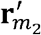 moves on the line crossing 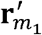 and normal to the calibration plane (Supplementary Fig. 9g and h). In the first case, we let 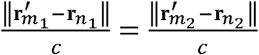 and observe how the probability of |**r**_*n*_1__ – **r**_*n*_2__| being small is violated. Because 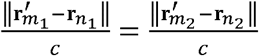 and locations 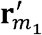 and 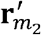 are with the same distance to the calibration plane, locations **r**_*n*_1__ and **r**_n_2__ must be on two circles with the same radius and their centers separated by 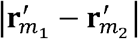. If 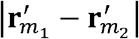 is small, for any **r**_*n*_1__ on one circle, there exists a **r**_*n*_2__ on the other circle such that |**r**_*n*_1__ – **r**_*n*_2__| is small, as shown in Supplementary Fig. 9e. If 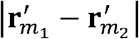 is large, a small value of |**r**_*n*_1__ – **r**_*n*_2__| requires both **r**_*n*_1__ and **r**_*n*_2__ to be close to the same intersection of the two circles, as depicted in Supplementary Fig. 9f Thus, the higher the value of 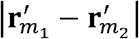, the lower the probability of |**r**_*n*_1__ – **r**_*n*_2__| being small. In summary, for the first case, the decay speed of 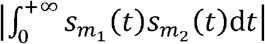 as 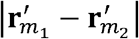 increases is mainly determined by MCC_*n*_1_, *n*_2__’s decay speed as |**r**_*n*_1__ – **r**_*n*_2__| increases. In the second case, we let *n*_1_ = *n*_2_ = *n* and observe the change of 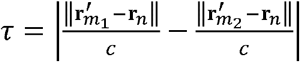. As 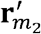 moves away from 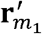 (from Supplementary Fig. 9g to h), 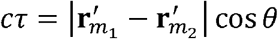 increases. Here, *θ* is the angle between vectors 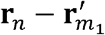 and 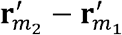. If 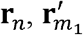, and 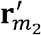 are on the same line, the calibrated virtual transducer at **r**_*n*_ is the most sensitive to the signals from 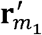 and 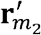 and cosθ ≈ 1. Thus, for the second case, the decay of 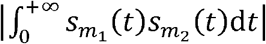 as 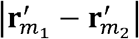 increases is mainly determined by the decay of 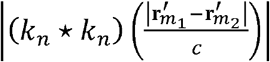. We compare the decay speed of MCC_*n*_1_,*n*_2__ as |**r**_*n*_1__ – **r**_*n*_2__| increases and that of 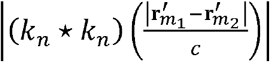 as 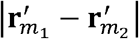 increases by observing the full-width values in Supplementary Fig. 9c and d. For both calibrated virtual transducers and all choices of *β*, values of FW_*L*_1_,*θ*_ and FW_*L*_2_,*θ*_ are of 4 to 8 times the value of FW_AC,β_.

The significant decay-speed difference between FW_*L*_1_,*θ*_ or FW_*L*_2_,*θ*_ with FW_*AC,θ*_ leads to the anisotropy of 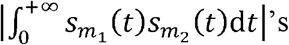 decay speed as 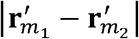 increases, which contributes to the anisotropy of the image resolution. It needs to be noted that, we have simplified the above analysis to identify the cross-correlation-related and autocorrelation-related limiting factors. In practice, the anisotropy is affected by both the maximum-cross-correlation matrix MCC_*n*_1_,*n*_2__ and the autocorrelation function (*k_n_* * *k_n_*)(*τ*) for all values of *n*_1_, *n*_2_, and *n* with different weights. These factors are incorporated into the iterative method and lead to the final anisotropy of resolution in the reconstructed image.

**Supplementary Fig. 1.**
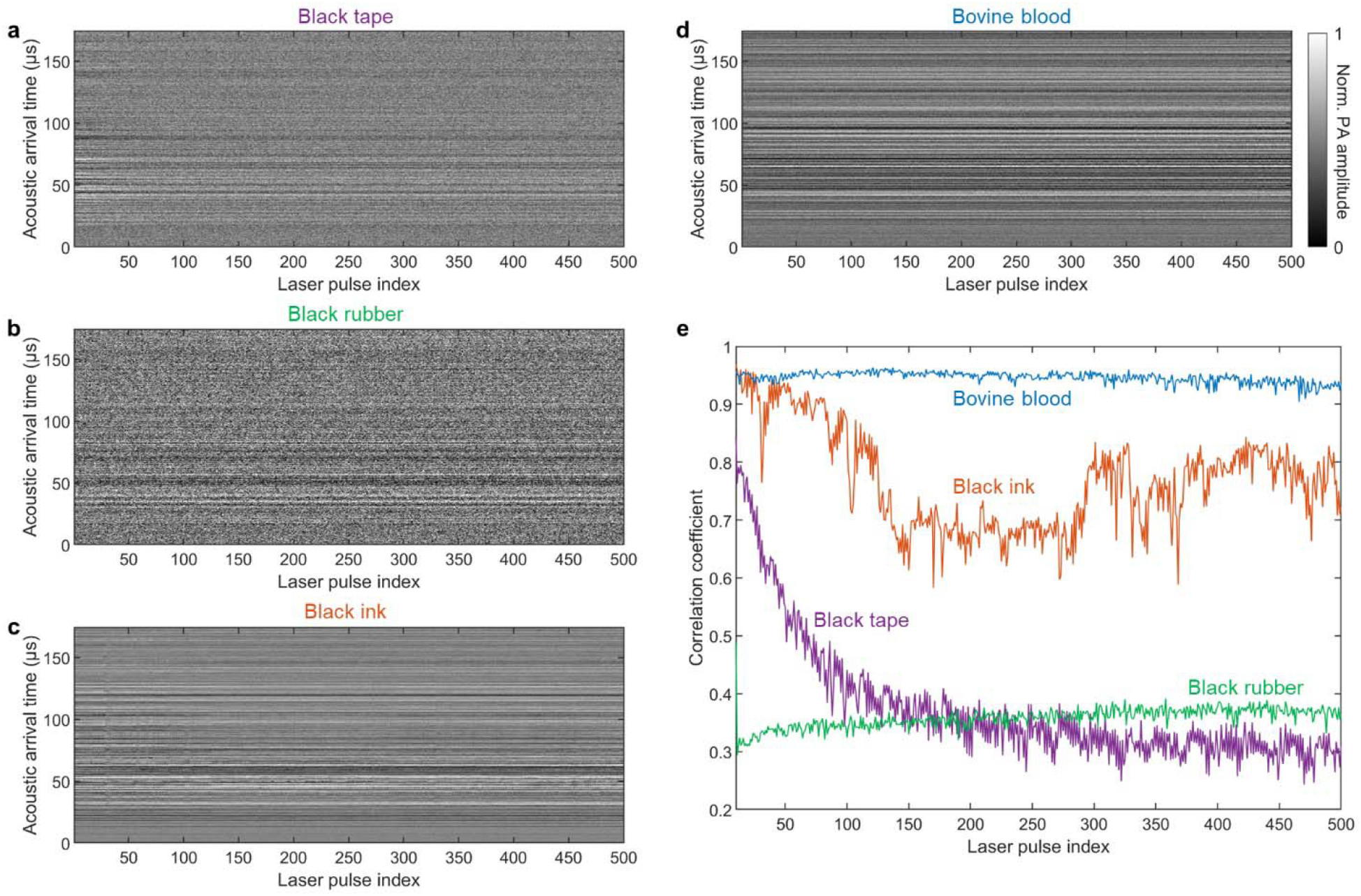
Comparison of uniform optical absorbers for calibration. **a–d**, PACTER signals generated by 500 consecutive laser pulses (8.6 μJ pulse energy, 1 kHz repetition rate) using black tape (**a**), black rubber (**b**), black ink (**c**), and bovine blood (**d**) as targets. Norm., normalized. **e**, Correlation coefficients between the PACTER signals and the average of the signals from the first 10 laser pulses.

**Supplementary Fig. 2.**
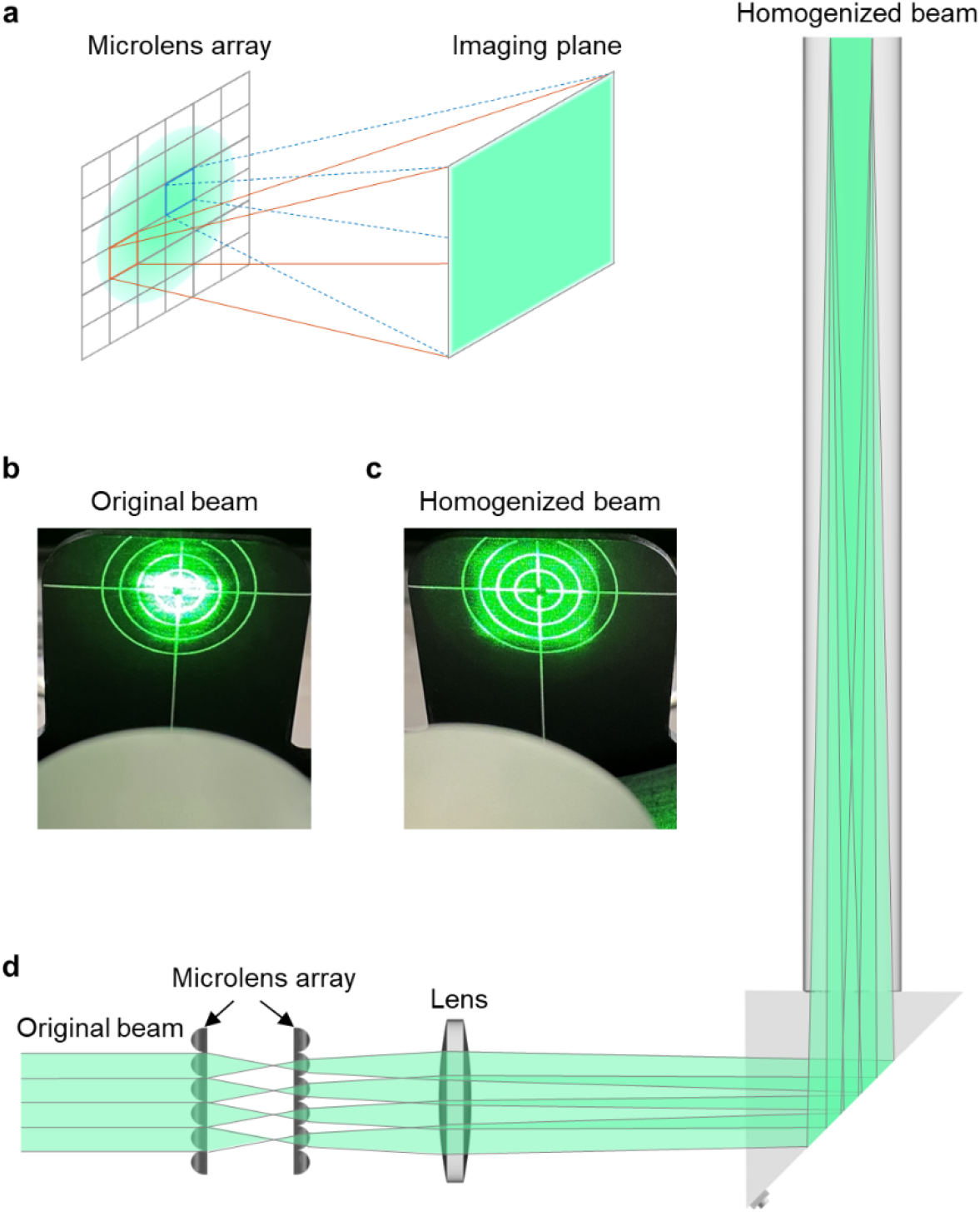
Fly’s eye homogenizer in PACTER. **a**, Schematic illustrating the working principle of the fly’s eye homogenizer. **b,c**, Photographs of the original beam from the laser (**b**) and the homogenized beam (**c**), captured using a laser alignment plate 1 cm away from the top of the ER. **d**, Schematic showing the optical path of the homogenizer and the ER.

**Supplementary Fig. 3.**
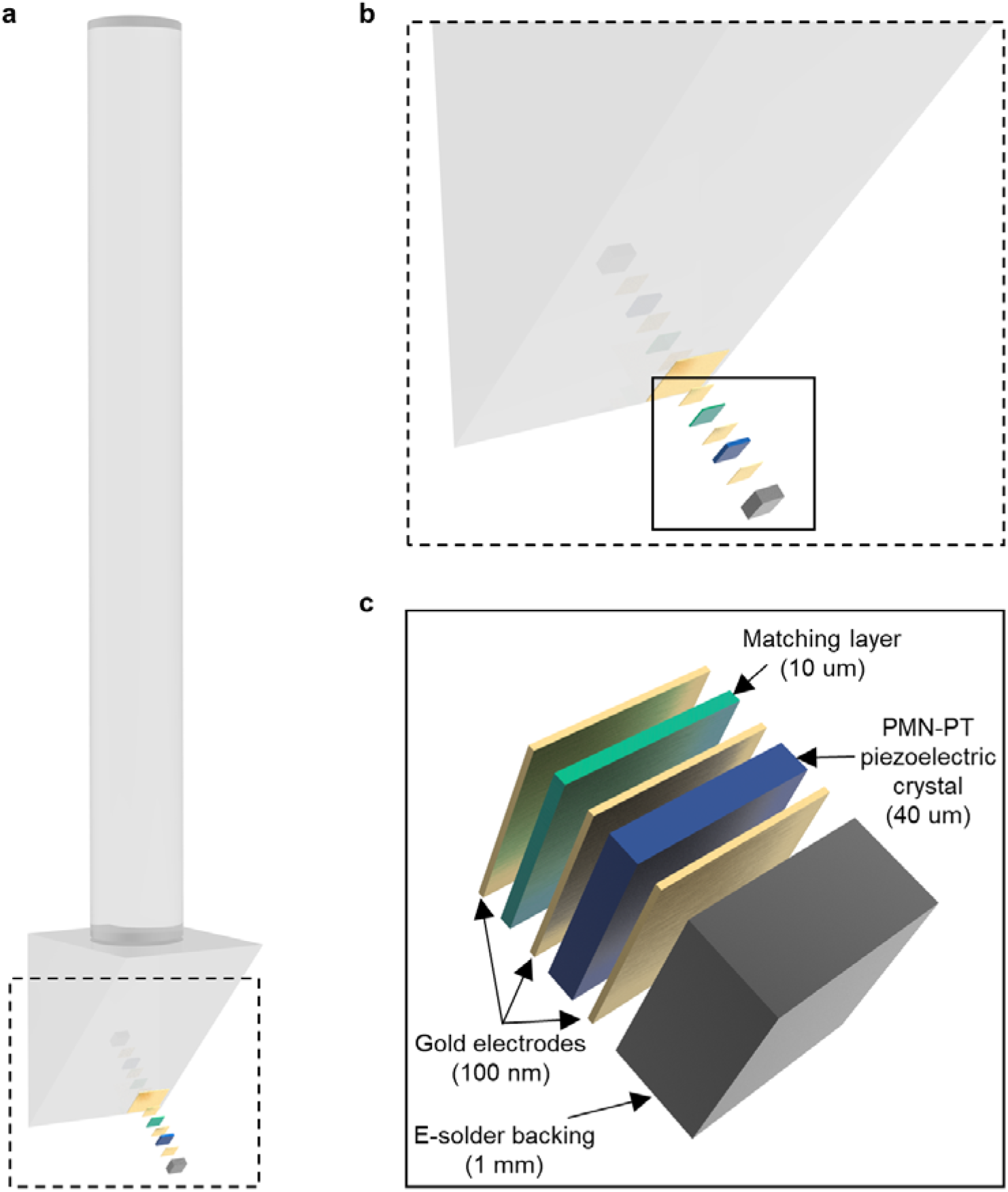
Fabrication of the single-element ultrasonic transducer. **a**, Schematic showing the ultrasonic transducer fabricated on the ER. **b**, Zoomed-in view of the black dashed box in **a**. **c**, Zoomed-in view of the black solid box in **b**. Thicknesses of different layers of materials are provided in brackets.

**Supplementary Fig. 4.**
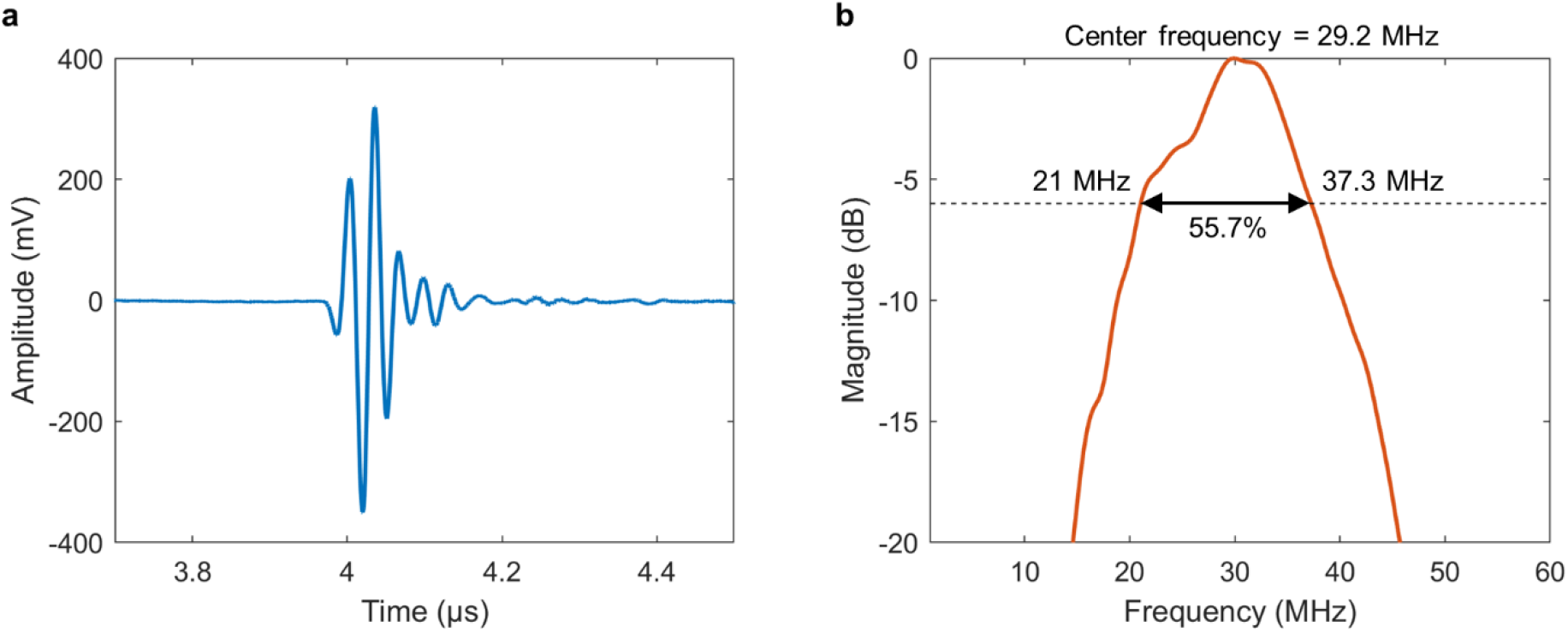
Characterization of the single-element ultrasonic transducer. Pulse-echo waveform (**a**) and −6 dB bandwidth (**b**) of the fabricated ultrasonic transducer measured before being glued to the ER.

**Supplementary Fig. 5.**
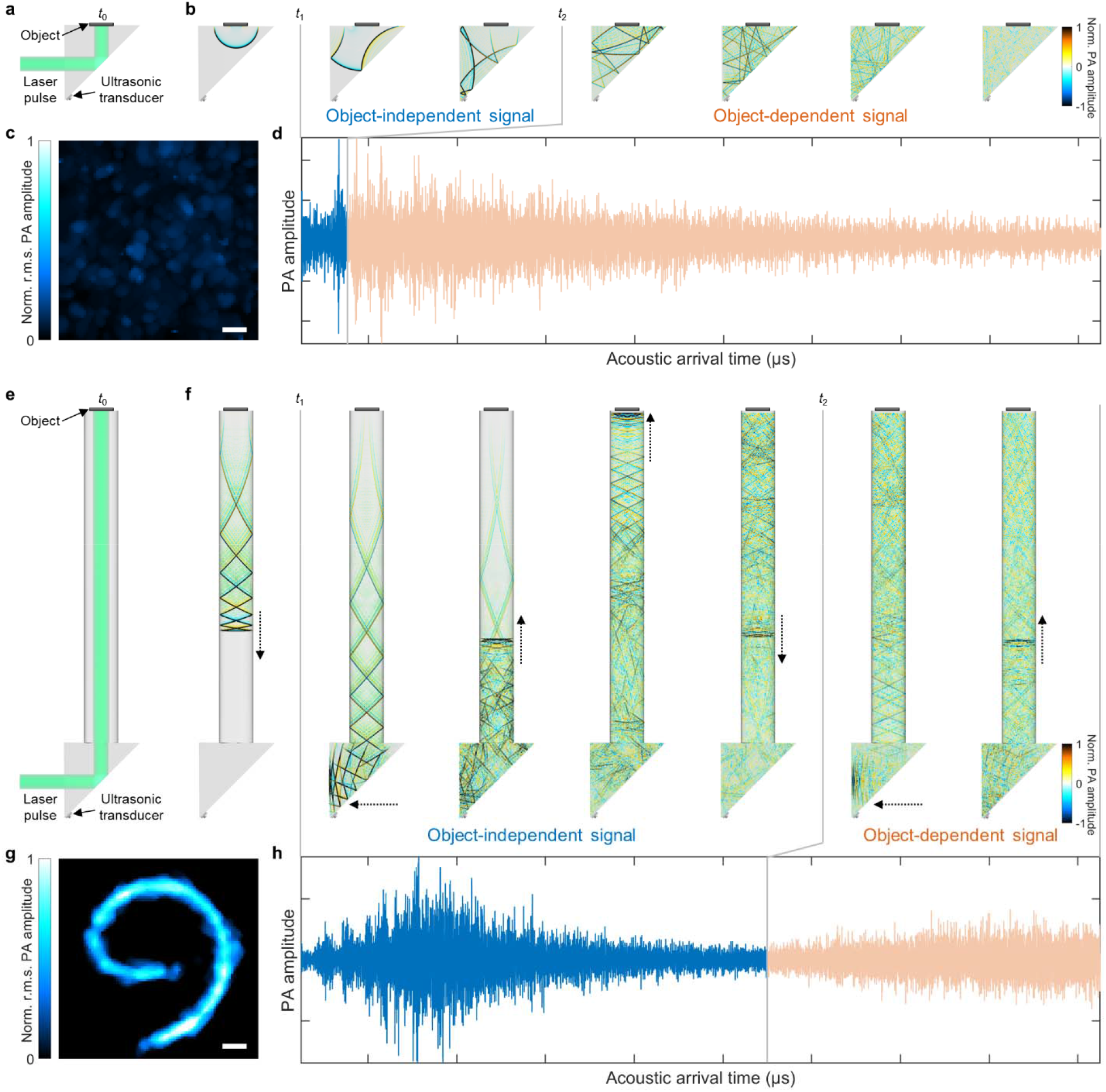
Object-dependent and -independent calibrations in PATER and PACTER, respectively. **a**, Optical path of the ER in PATER. *t*_0_ denotes the time instance of laser pulse illumination. **b**, Time-lapse simulation of the PA signal propagating in the ER in PATER. *t*_1_ denotes the time instance when the transducer starts to detect the PA signal. *t*_2_ denotes the time instance when the transducer starts to detect the object-dependent PA signal, which is reflected from the object and affected by the boundary condition. Norm., normalized. **c**, Reconstructed image of a black wire using the ER in PATER calibrated with bovine blood, showing the object dependence of calibration. **d**, PA signal detected by a transducer attached to the ER in PATER. **e**, Optical path of the ER in PACTER. **f**, Time-lapse simulation of the PA signal propagating in the ER in PACTER. Black dotted arrows denote the propagating direction of the acoustic wavefront. The definitions of *t*_0_, *t*_1_, *t*_2_ are identical to those in **a** and **b**. **g**, Reconstructed image of a black wire using the ER in PACTER calibrated with bovine blood, showing the object independence or universality of calibration. **h**, PA signal detected by the transducer fabricated on the ER in PACTER. Object-independent and -dependent signal segments are color-coded in blue and light orange, respectively. Scale bars, 1 mm.

**Supplementary Fig. 6.**
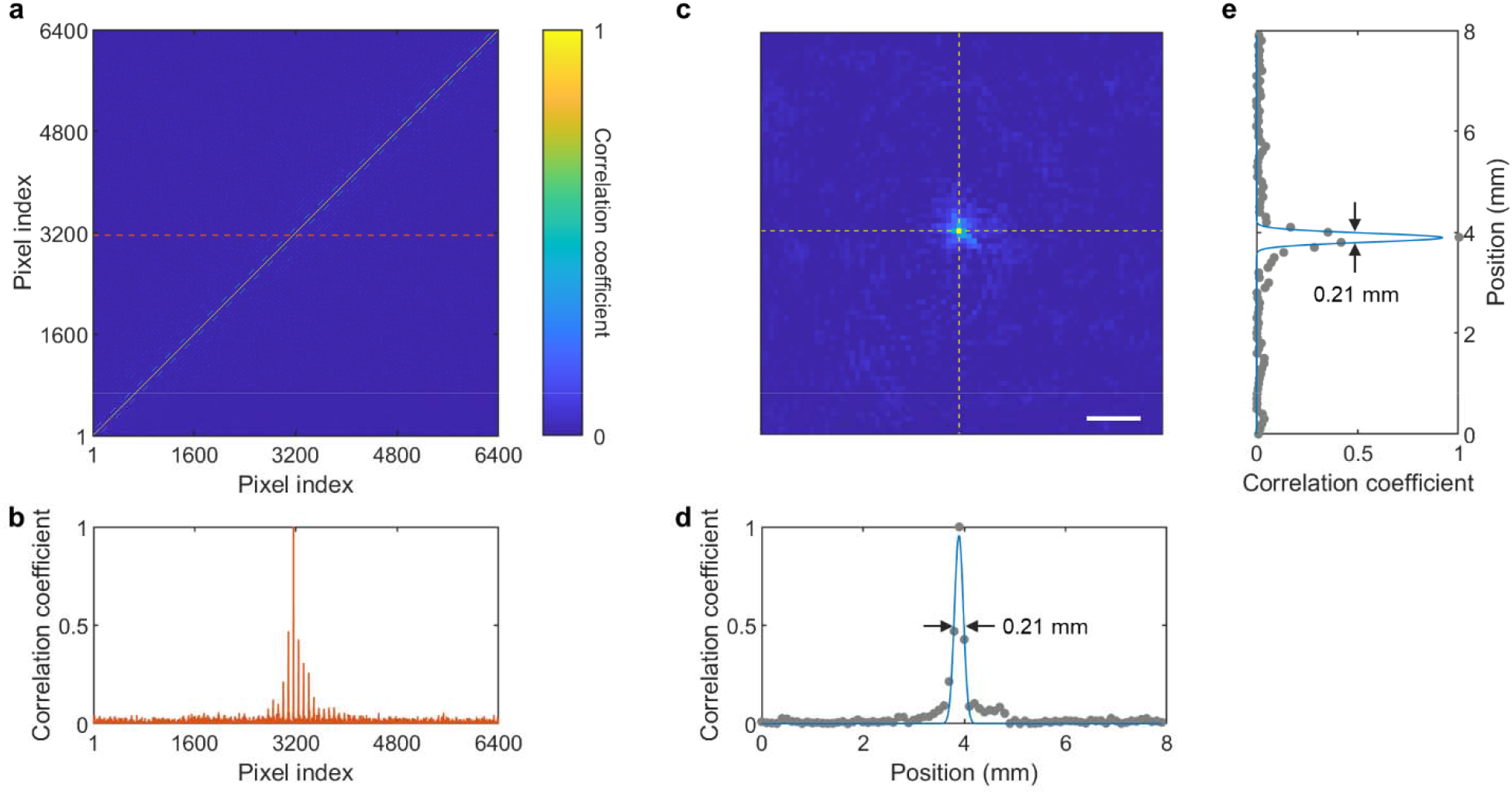
PACTER signals from calibration using bovine blood. **a**, Cross-correlation map of the PACTER signals from the calibration pixels (80 by 80 steps with a 0.1 mm step size), where the diagonal line represents autocorrelation. **b**, Line profile along the orange dashed line in **a**. **c**, Cross-correlation map (80 by 80 pixels) reshaped from the crosscorrelation profile in **b**. Scale bar, 1 mm. **d**,**e**, Line profiles along the horizontal (**d**) and vertical (**e**) yellow dashed lines in **c**. Both FWHMs of the fitted profiles are ~0.21 mm.

**Supplementary Fig. 7.**
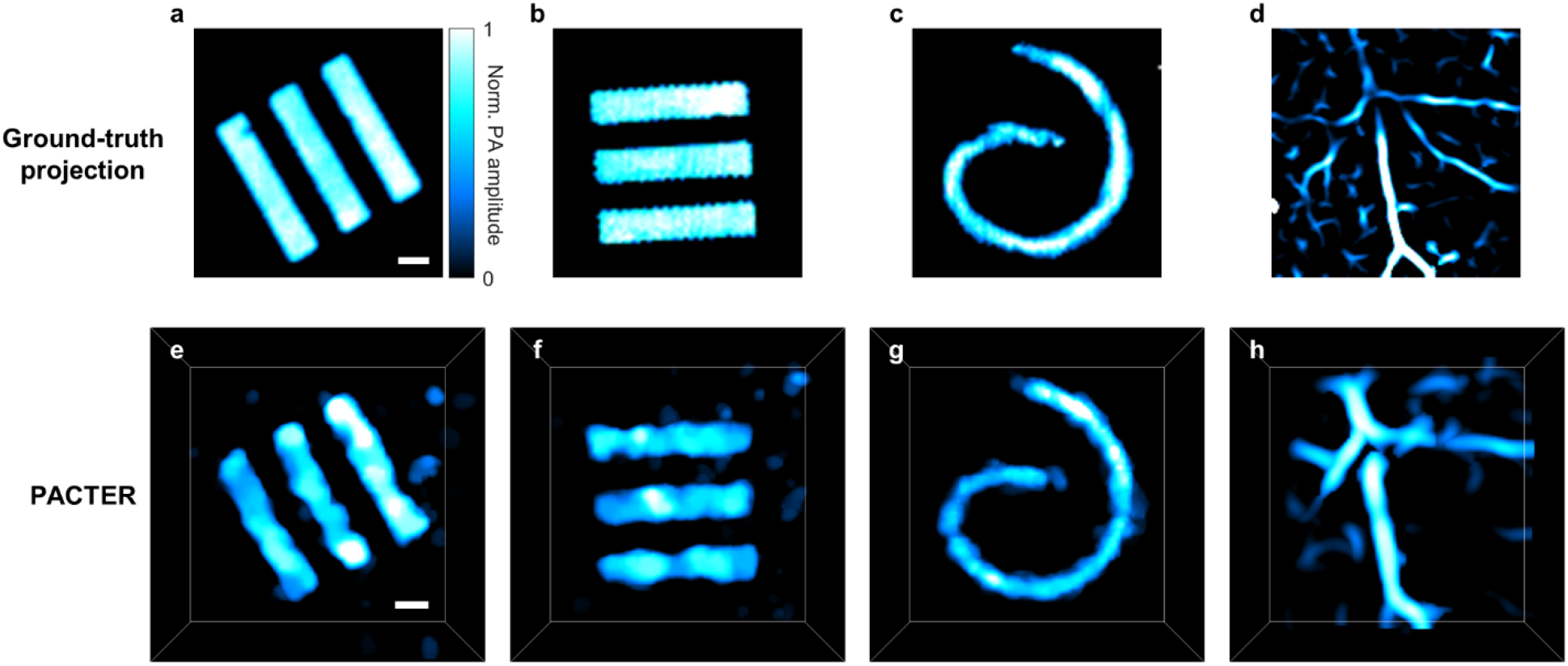
Validation of PACTER reconstruction. **a-d**, Ground-truth projection images of the objects: bars printed with black ink on a transparent film (**a,b**), curved black wire (**c**), and mouse abdominal vasculature *in vivo* (**d**). The images were acquired in a way similar to the calibration procedure, except that the bovine blood was replaced by the objects. The images were formed by the root-mean-squared projections of the PACTER signals as the focused laser beam scanned across the objects. Norm., normalized. **e–h**, Perspective views of 3D PACTER images of the objects in **a–d** acquired through the imaging procedure using a homogenized beam. Scale bars, 1 mm.

**Supplementary Fig. 8.**
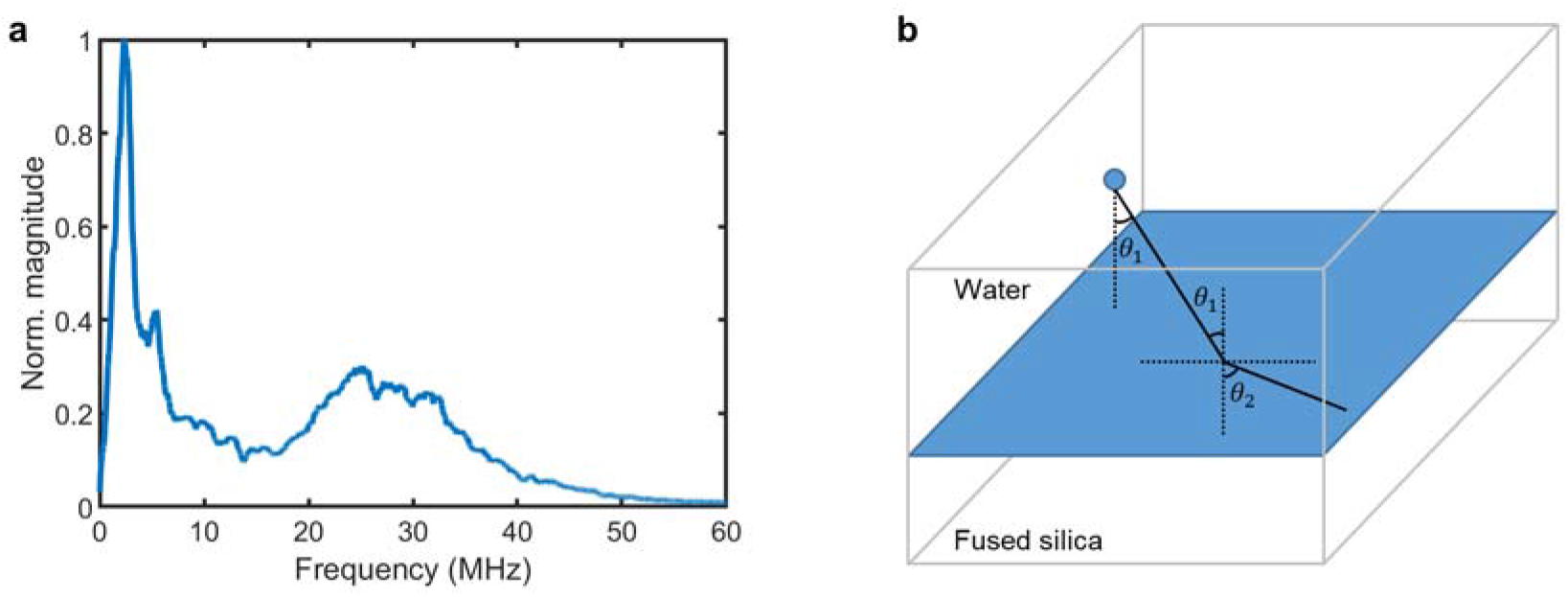
Spatial resolution analysis from the virtual-transducer-array perspective. **a**Spectrum of the PACTER signal from the phantom shown in Fig. 3f. **b** Refraction of the ultrasonic wave at the boundary between water and fused silica. *θ*_1_ is the incident angle. *θ*_2_ is the refraction angle.

**Supplementary Fig. 9.**
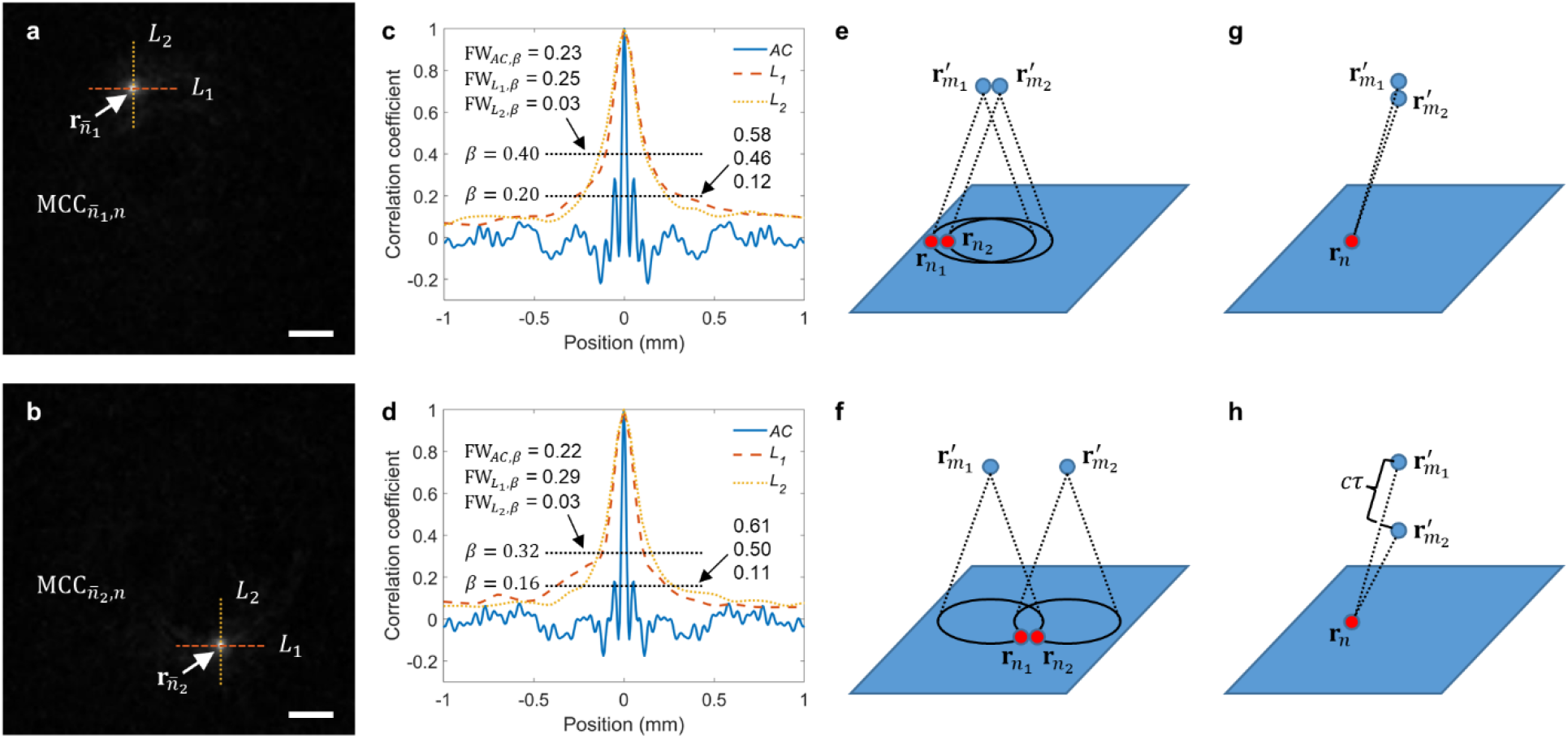
Spatial resolution analysis from the system-matrix perspective. **a-b**, Two slices of the maximum-cross-correlation matrix (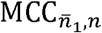 and 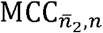, each value means the maximum cross correlation between two impulse responses in the calibration) corresponding to the 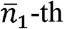 and 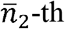 calibrated virtual transducers, respectively. Two lines *L*_1_ and *L*_2_ are drawn in each image. Scale bars, 1 mm. **c-d**, Maximum-cross-correlation values along lines *L*_1_ and *L*_2_ aligned with the values of 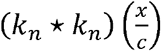 for the 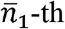 and 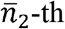 calibrated virtual transducers, respectively. **e-f**, A source point at 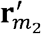 moves away from another source point at 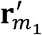 (from **e** to **f**) with both source points on a line parallel to the calibration plane. In each image, we mark two close calibrated virtual transducers **r**_*n*_1__ and **r**_*n*_2__ which satisfy 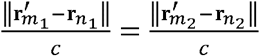. **g-h**, A source point at 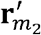 moves away from another source point at 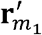 (from **g** to **h**) with both source points on a line normal to the calibration plane. The same calibrated virtual transducer **r**_*n*_ is marked in both **g** and **h** with the difference between its distances to 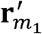 and 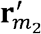 denoted as *cτ*.

**Supplementary Fig. 10.**
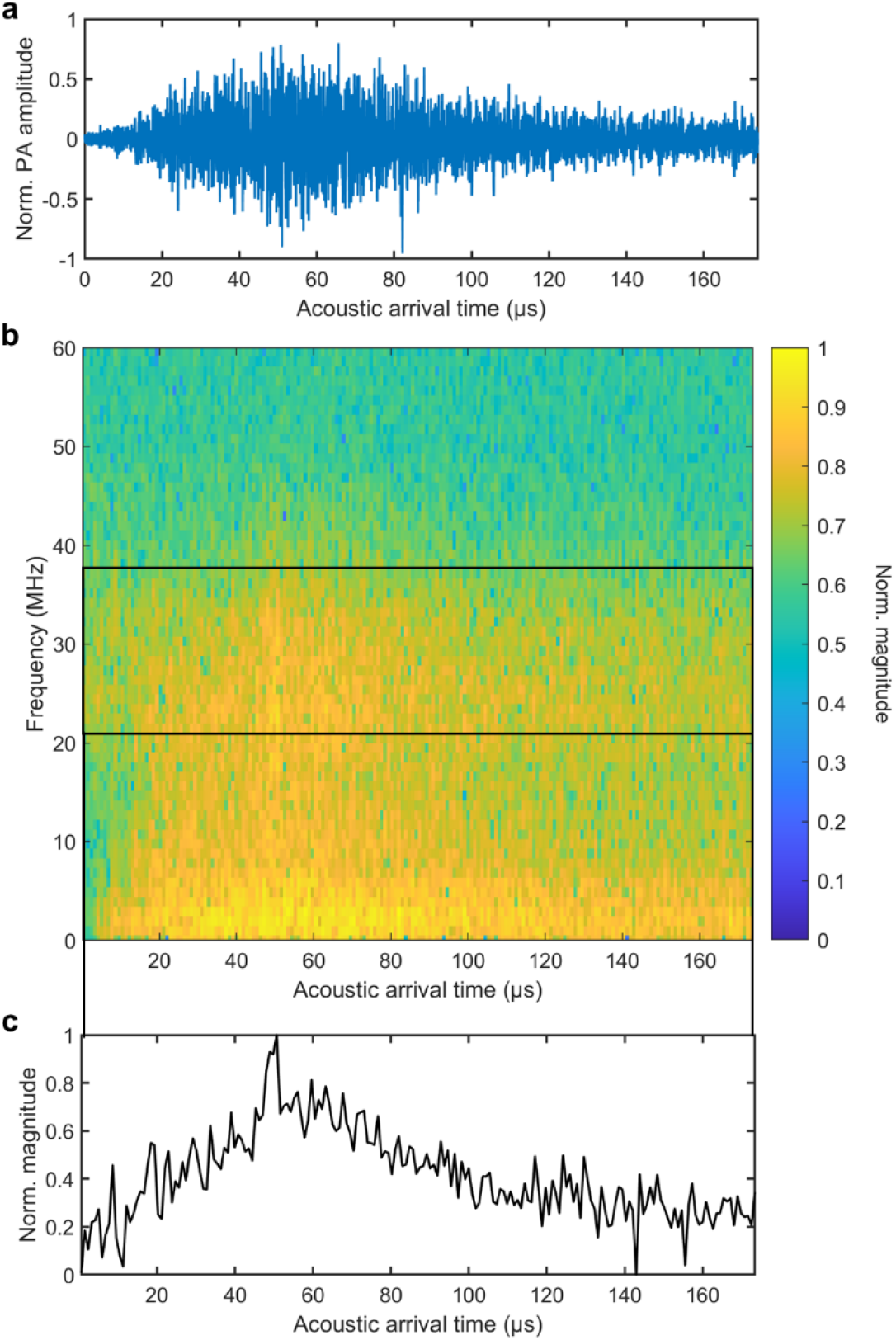
Spectrogram of the PACTER signal. **a**, PACTER signal of bovine blood in the time domain. Norm., normalized. **b**, Spectrogram of the PACTER signal in **a** calculated using a short-time Fourier transformation (256 samples per section, 32 samples overlapped between sections). **c**, Spectral magnitude averaged over the bandwidth of the ultrasonic transducer, i.e., from 21 MHz to 37.3 MHz, versus time, corresponding to the black box in **b**.

**Supplementary Fig. 11.**
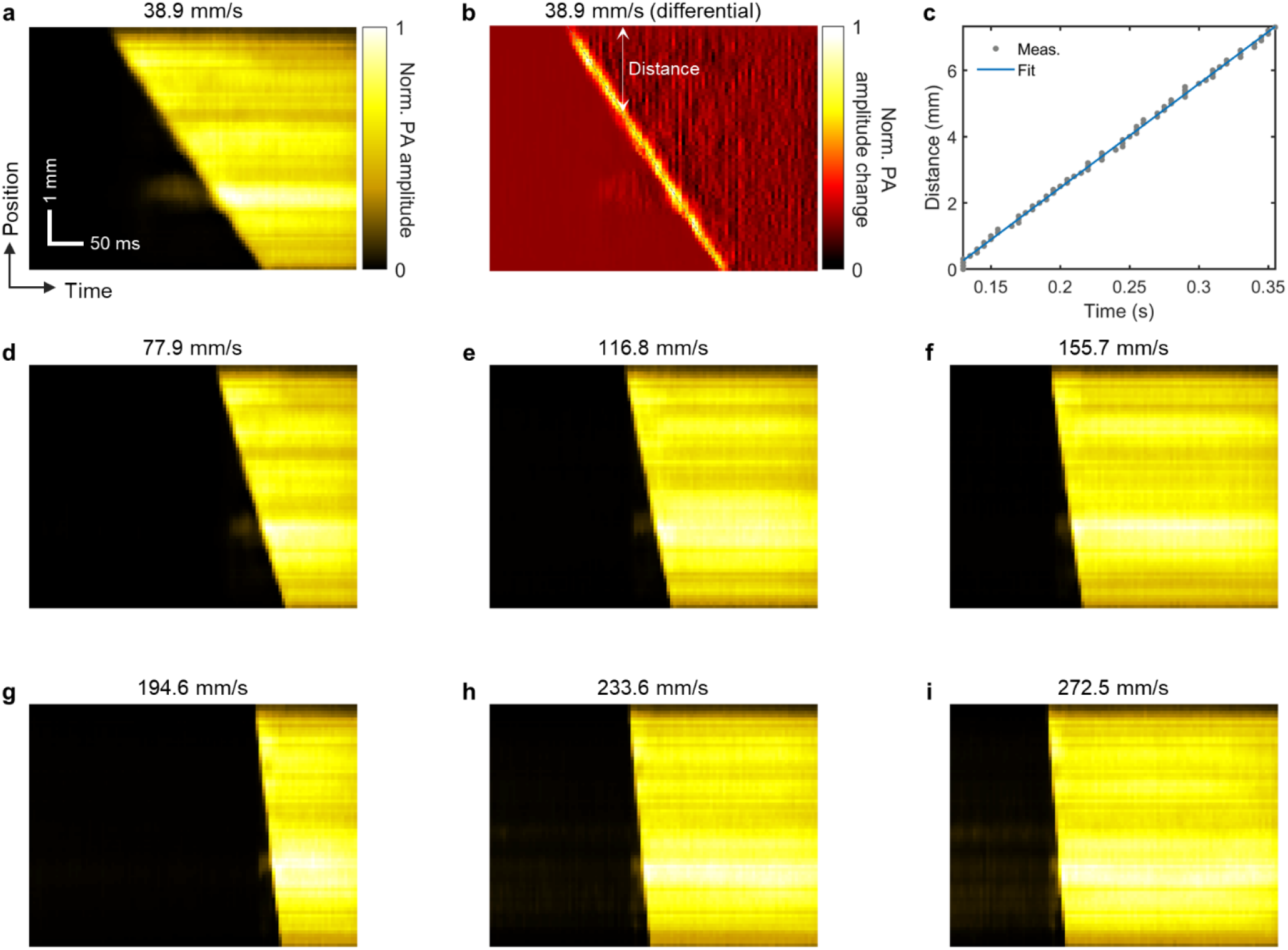
PACTER of bovine blood flushing through a tube. **a**, PA amplitudes along the tube (1D image) flushed by the blood with a speed of 38.9 mm/s versus time. Norm., normalized. **b**, Differential image of **a** (difference between adjacent voxels along the tube in **a**versus time) showing the distance the blood front travels over time. **c**, Traveling distance of the blood front plotted over time. Blue curve represents a linear fit. **d–i**, PA amplitudes along the tube (1D images) flushed by the blood with different speeds versus time.

**Supplementary Fig. 12.**
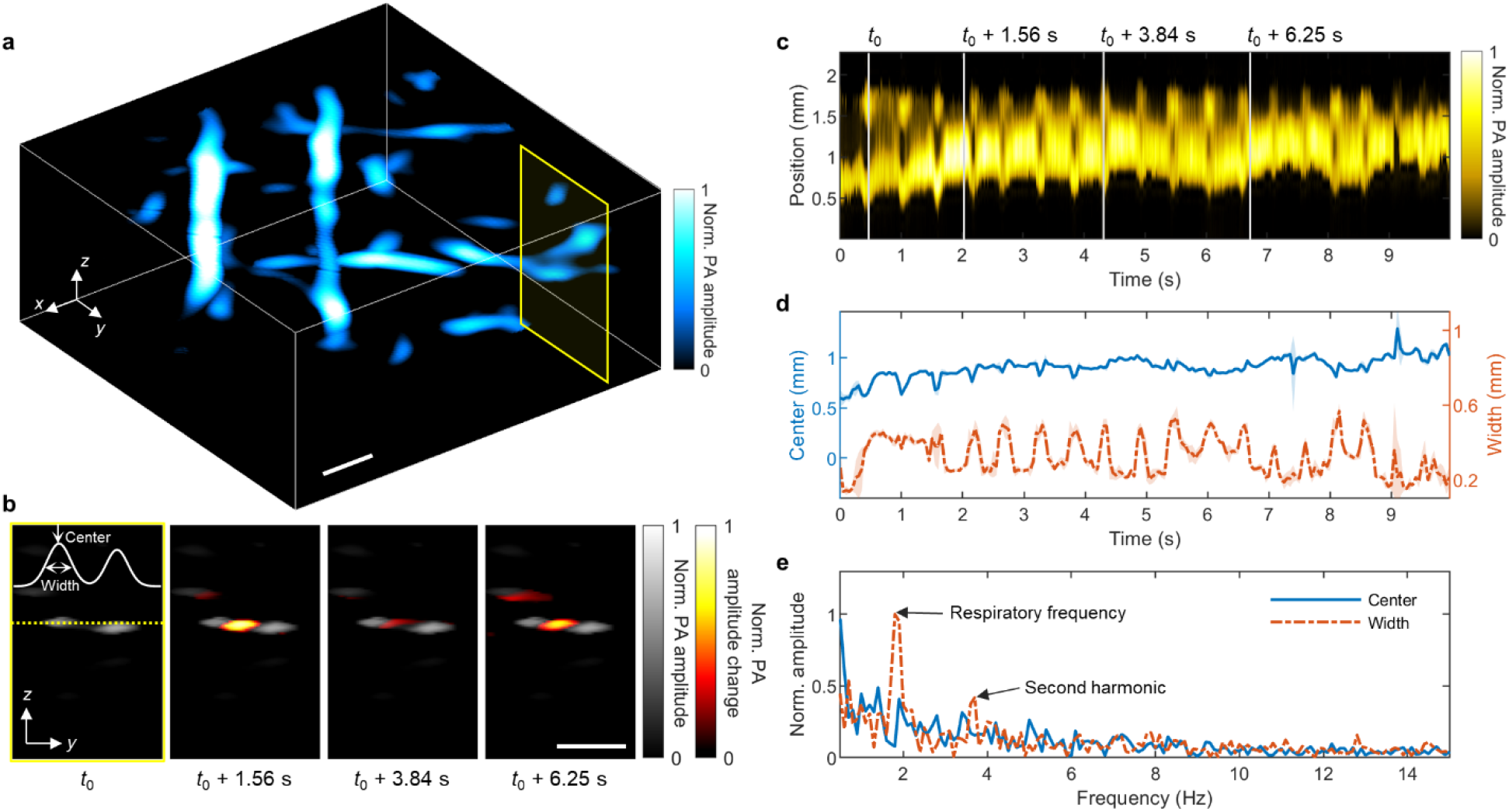
PACTER of mouse hemodynamics *in vivo*. **a**, 3D PACTER image of the abdominal vasculature of mouse 3. Norm., normalized. **b**, Crosssectional 2D images corresponding to the yellow rectangle in **a** at four different time instances from the 4D PACTER datasets. *t*_0_ = 0.49 s. The white solid curve represents a two-term Gaussian fit of the vessels’ profile denoted by the yellow dashed line. Differences from the first image are highlighted in color. **c**, PA amplitudes along the yellow dashed line (1D image) in **b**versus time, where the time instances in **b** are labeled with vertical gray lines. **d**, Blue solid and orange dash-dotted curves represent the center positions and widths of the vessel on the left (based on the first term of the Gaussian fit in **b**) versus time. Shaded areas denote the standard deviations (*n* = 5). **e**, Fourier transforms of the center positions and widths of the vessel in **d**, showing the respiratory frequency from the vessel widths only. Scale bars, 1 mm.

**Supplementary Fig. 13.**
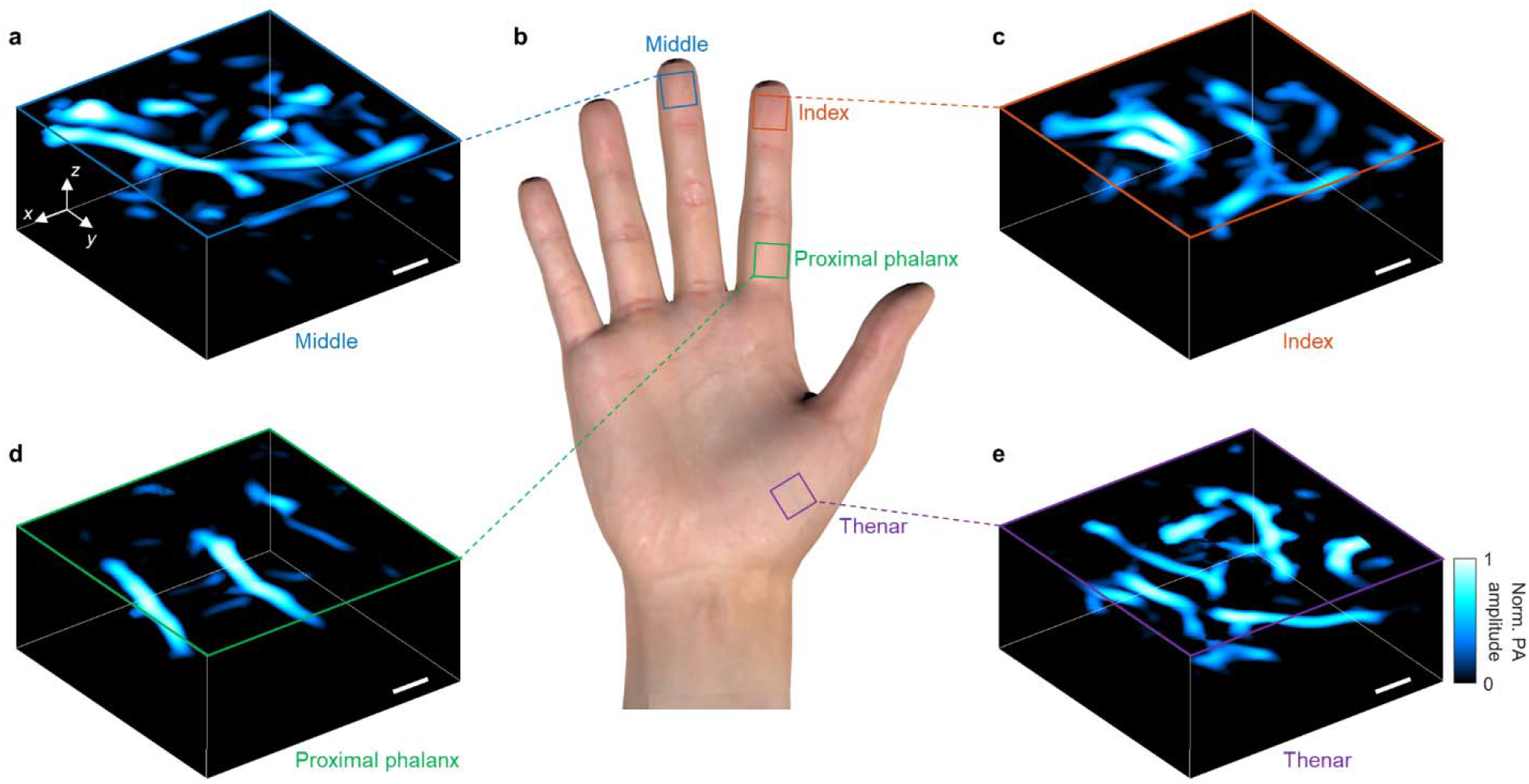
PACTER of human hand vasculature *in vivo*. **a,c–e**, 3D PACTER images of the vasculature in a middle finger (**a**), an index finger (**c**), a proximal phalanx region (**d**), and a thenar region (**e**). Norm., normalized. **b**, Photograph of a human hand showing the imaged regions. Scale bars, 1 mm.

**Supplementary Fig. 14.**
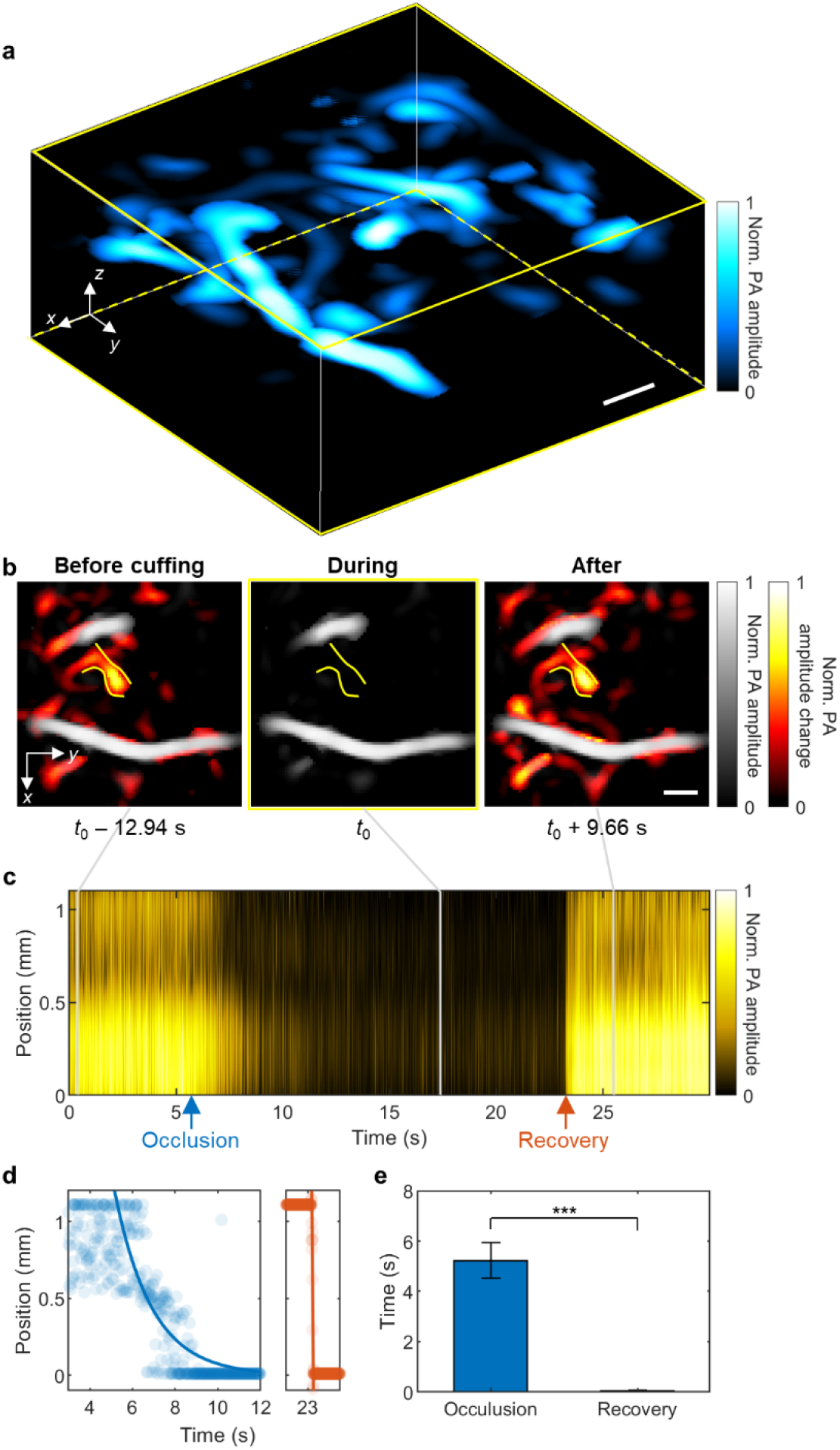
PACTER of human hand hemodynamics *in vivo*. **a**, 3D PACTER image of the thenar vasculature of participant 1, in a region different from that in Fig. 5**b**. Norm., normalized. **b**, Maximum amplitude projections of the 3D volumes from the 4D PACTER datasets along the *z* axis in **a** at the time instances before, during, and after cuffing. *t*_0_ = 15.34 s. The solid lines flank the vessel under investigation. Differences from the *t*_0_ image are highlighted in color. **c**, PA amplitudes along the vessel (1D image) in **b** versus time, where the time instances in **b** are labeled with vertical gray lines. The blue and orange arrows indicate peak responses in the occlusion and recovery phases, respectively. **d**, Positions of the blood front along the blood vessel during the occlusion (left) and recovery (right) phases in **c**. The left blue curve is an exponential fit with an occlusion rate of 0.6 ± 0.1 m/s, whereas the right orange curve is a linear fit showing the blood flow speed of 22.4 ± 6.4 m/s. **e**, Comparison between the durations of the occlusion and recovery phases in **d**. \*\*\**P* < 0.001, calculated by the two-sample *t*-test. Scale bars, 1 mm.

**Supplementary Fig. 15.**
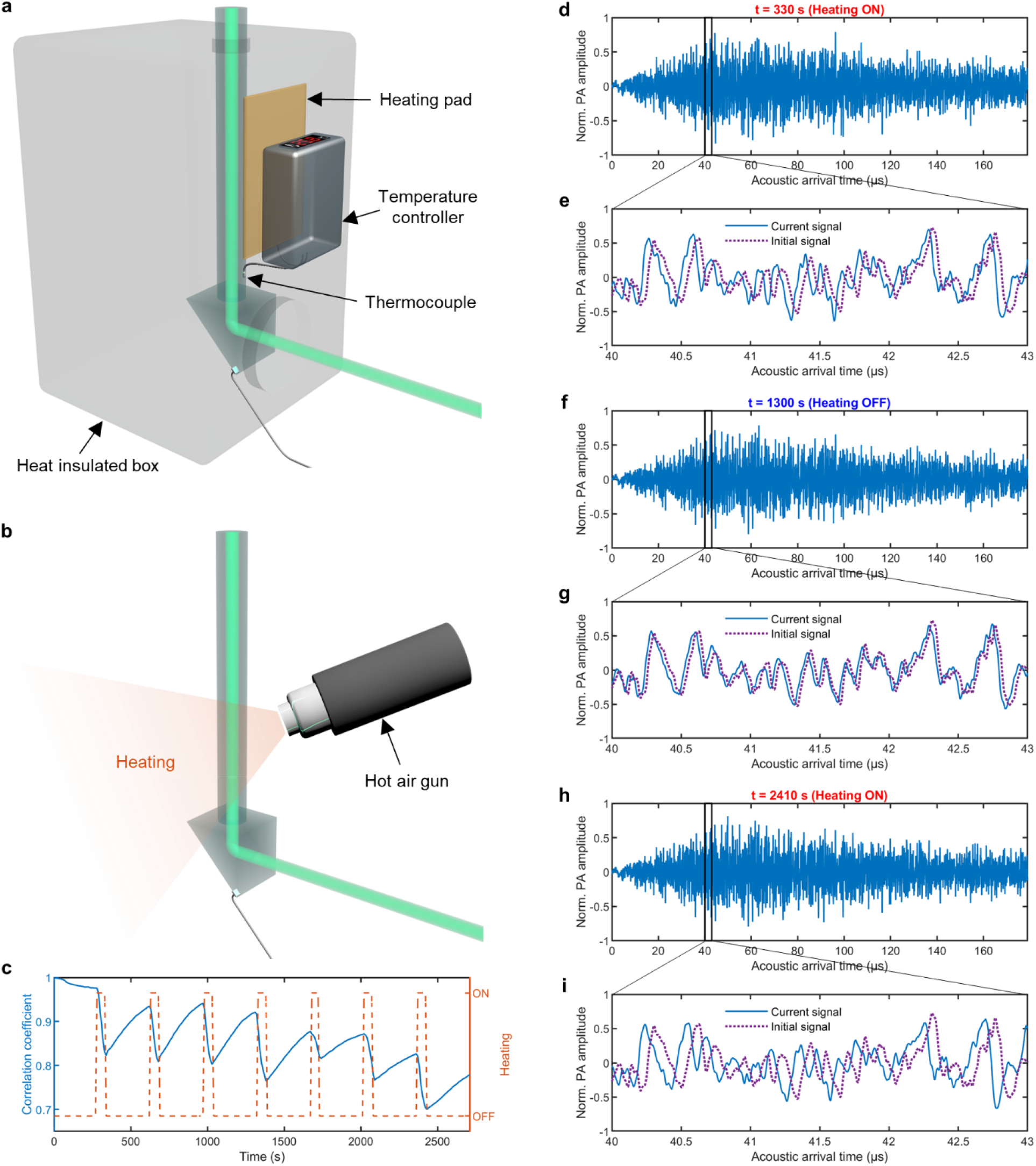
Temperature stabilization in PACTER. **a**, Schematic of the temperature stabilizing box in the PACTER system. **b**, Illustration of the experiment showing the temperature dependence of the PACTER system. During acquisition of PACTER signals, the hot air gun is turned ON and OFF to provide periodic heating to the ER. **c**, Correlation coefficient (blue solid curve) between each PACTER signal and the initial signal acquired using bovine blood, and the ON and OFF status (orange dashed curve) of the hot air gun. **d,f,h**, PACTER signals acquired at 330 s with heating on (**d**), 1300 s with heating off (**f**), and 2410 s with heating on (**h**). Norm., normalized. **e,g,i**, The blue solid lines represent the zoomed-in views of the black boxes in **d** (**e**), **f** (**g**), and **h** (**i**). The purple dotted lines show the initial signal acquired at 0 s.

## Description of Additional Supplementary Files

**Supplementary Video 1**

Principle and implementation of PACTER (with narration).

**Supplementary Video 2**

Simulation of PA wave propagation in the ERs in PATER (left) and PACTER (right) and the PA signals detected by the ultrasonic transducers attached to or fabricated on the cavities. The object-independent signal over a sufficient duration in PACTER enables object-independent universal calibration. Norm., normalized.

**Supplementary Video 3**

4D (3D in space and 1D in time) PACTER image and its maximum amplitude projection of bovine blood flushing through an S-shaped tube. Norm., normalized.

**Supplementary Video 4**

4D PACTER images of bovine blood flowing through a tube at different speeds. Norm., normalized.

**Supplementary Video 5**

4D PACTER image and its maximum amplitude projection of bovine blood flowing through a tube with a speed of 272.5 mm/s, captured at 1,000 volumes per second (i.e., 1 ms per volume). Norm., normalized.

**Supplementary Video 6**

Top-left, 4D *in vivo* PACTER image of the abdominal vasculature of mouse 1. Norm., normalized. Top-right, 3D (2D in space and 1D in time) cross-sectional image corresponding to the yellow rectangle. Middle, PA amplitudes along the yellow dashed line (1D image) versus time. Bottom, center positions (blue solid curve) and widths (orange dash-dotted curve) of the vessel profiled by the yellow dashed line versus time. The shaded areas denote the standard deviations (*n* = 5).

**Supplementary Video 7**

Top-left, 4D *in vivo* PACTER image of the abdominal vasculature of mouse 2. Norm., normalized. Top-right, 3D (2D in space and 1D in time) cross-sectional image corresponding to the magenta rectangle. Middle, PA amplitudes along the magenta dashed line (1D image) versus time. Bottom, center positions (blue solid curve) and widths (orange dash-dotted curve) of the vessel profiled by the magenta dashed line versus time. The shaded areas denote the standard deviations (*n* = 5).

**Supplementary Video 8**

Top-left, 4D *in vivo* PACTER image of the abdominal vasculature of mouse 3. Norm., normalized. Top-right, 3D (2D in space and 1D in time) cross-sectional image corresponding to the yellow rectangle. Middle, PA amplitudes along the yellow dashed line (1D image) versus time. Bottom, center positions (blue solid curve) and widths (orange dash-dotted curve) of the vessel profiled by the yellow dashed line versus time. The shaded areas denote the standard deviations (*n* = 5).

**Supplementary Video 9**

Top-left, 4D *in vivo* PACTER image of the thenar vasculature of participant 1. Norm., normalized. Top-right, maximum amplitude projection of the 3D volume. Middle, PA amplitudes along the yellow dashed line (1D image) versus time. Bottom, total PA amplitude (sum of all voxel values along the dashed line) versus time. The shaded areas denote the standard deviations (*n* = 5). The orange and cyan dashed vertical lines indicate the start times of occlusion and recovery, respectively.

**Supplementary Video 10**

Top-left, 4D *in vivo* PACTER image of the thenar vasculature of participant 2. Norm., normalized. Top-right, maximum amplitude projection of the 3D volume. Middle, PA amplitudes along the magenta dashed line (1D image) versus time. Bottom, total PA amplitude (sum of all voxel values along the dashed line) versus time. The shaded areas denote the standard deviations (*n* = 5). The orange and cyan dashed vertical lines indicate the start times of occlusion and recovery, respectively.

**Supplementary Video 11**

Top-left, 4D *in vivo* PACTER image of the thenar vasculature of participant 1, in a region different from that in Supplementary Video 9. Norm., normalized. Top-right, maximum amplitude projection of the 3D volume. Middle, PA amplitudes along the yellow dashed line (1D image) versus time. Bottom, total PA amplitude (sum of all voxel values along the dashed line) versus time. The shaded areas denote the standard deviations (*n*= 5). The orange and cyan dashed vertical lines indicate the start times of occlusion and recovery, respectively.

**Supplementary Video 12**

Top, PACTER signals acquired using bovine blood while the ER was periodically heated by a hot air gun. Middle, zoomed-in view of the signals in the black box in relation to the initial signal acquired at 0 s shown as purple dotted line. Bottom, correlation coefficient between the PACTER signal and the initial signal versus time. The orange dashed curve denotes the ON and OFF status of the hot air gun.

